# From interaction networks to interfaces: Scanning intrinsically disordered regions using AlphaFold2

**DOI:** 10.1101/2023.05.25.542287

**Authors:** Hélène Bret, Jessica Andreani, Raphaël Guerois

## Abstract

The revolution brought about by AlphaFold2 and the performance of AlphaFold2-Multimer open promising perspectives to unravel the complexity of protein-protein interaction networks. Nevertheless, the analysis of interaction networks obtained from proteomics experiments does not systematically provide the delimitations of the interaction regions. This is of particular concern in the case of interactions mediated by intrinsically disordered regions, in which the interaction site is generally small. Using a dataset of protein-peptide complexes involving intrinsically disordered protein regions that are non-redundant with the structures used in AlphaFold2 training, we show that when using the full sequences of the proteins involved in the interaction networks, AlphaFold2-Multimer only achieves 40% success rate in identifying the correct site and structure of the interface. By delineating the interaction region into fragments of decreasing size and combining different strategies for integrating evolutionary information, we managed to raise this success rate up to 90%. Beyond the correct identification of the interaction site, our study also explores specificity issues. We show the advantages and limitations of using the AlphaFold2 confidence score to discriminate between alternative binding partners, a task that can be particularly challenging in the case of small interaction motifs.

## Introduction

Protein interactions are crucial for a vast number of processes in living organisms. Strong evidence points to the biological importance of interactions mediated by intrinsically disordered protein regions (IDRs), such as short linear motifs, in particular for regulation, transport and signaling, and in a number of human pathologies ^1, 2, 3^. Established resources exist to identify already annotated binding motifs, such as the Eukaryotic Linear Motif (ELM) repository ^4^, to visualize evolutionary properties ^5^ and to screen full protein sequences for disordered stretches that might fold upon binding, as with the IUPred server ^6^.

Protein interactions are connected within complex networks called interactomes, which can be derived from large amounts of experimental data such as proteomics. Much effort has been invested into mapping and modeling interactions at the scale of these interactomes ^7, 8^. In these networks, most protein-protein interactions evolve under negative selection to maintain function and many of them can rewire ^9^, although at different evolutionary rates: stable protein complexes evolve more slowly than most domain-motif interactions ^10^. Interactions in evolutionarily old, housekeeping protein complexes are conserved across different contexts (cell types, tissues and conditions) while evolutionarily young interactions and those mediated by disordered regions are more versatile ^11, 12^. Evolutionary conservation has long been recognized as relevant to detect binding motifs in disordered regions, as reviewed in ^13^; however, the quality of the multiple sequence alignment (MSA) used for detection is particularly crucial ^14^.

AlphaFold2 revolutionized structure prediction for single proteins by leveraging deep learning approaches to extract signal from MSAs and output protein atomic 3D coordinates in an end-to-end manner ^15^. AlphaFold2 structure predictions for the entire human proteome ^16^ hinted that low prediction quality could pinpoint regions likely to be intrinsically disordered. Subsequent studies confirmed that AlphaFold2, although trained only on single proteins with a folded structure, can be used as an intrinsic disorder predictor by repurposing low-confidence residue predictions ^17, 18, 19^. AlphaFold2 low-confidence predictions on protein surfaces might also be indicative of possible binding regions ^20^.

Very soon after its release, AlphaFold2 was also tested for its capacity to predict protein-protein interactions. Despite not being designed for this purpose, AlphaFold2 outperformed traditional methods for the structural prediction of complexes between globular protein domains, in terms of both success rate and model quality ^21, 22, 23, 24, 25, 26, 27^. AlphaFold-Multimer, specifically retrained on protein complexes, displayed improved performance for interface modeling over the original AlphaFold2 ^21, 22, 28^. At a wider scale, a systematic exploration of the yeast interactome used prefiltering with a fast version of RoseTTAFold ^29^ followed by AlphaFold2 structure prediction ^30^. This opened exciting perspectives for the use of AlphaFold2 not only for complex structure prediction, but also as an *in silico* screening tool for interactions.

AlphaFold2 predictions are sensitive to the input MSA and protein delimitations. For instance, AlphaFold2 can be made to predict alternative conformational states for some proteins through manipulation of the MSA either by subsampling ^31^ or by *in silico* mutagenesis ^32^. For complexes, the generation of a paired MSA, where species are matched between homologs of the different protein partners, was not found to be necessary for AlphaFold2 to pick up interaction signal ^21, 25^, although combining unpaired and paired MSAs gave the best results ^21^. The AlphaPulldown package allows users to select or screen protein fragments for modeling protein complexes, since some interactions cannot be predicted if the full-length sequences are provided to AlphaFold2 ^33^.

Interactions mediated by short peptides within disordered protein regions are quite specific and thus require extra care for handling by AlphaFold2. Indeed, conformational versatility is even higher and covariation signal is weaker than for globular complexes ^34^. Traditional tools to predict protein-peptide interactions include mostly docking approaches, recently reviewed in ^35, 36^; some of these also make use of evolutionary information ^13^. Several recent studies have addressed the ability of AlphaFold2 to predict protein-peptide complexes. An early implementation already showed interesting predictive capacity, including in cases where the peptide induces a large conformational change of the protein and docking therefore most likely fails, and without the need for a peptide MSA ^37^. InterPepScore ^38^, a graph neural network used to score protein-peptide complexes for improving Rosetta FlexPepDock refinement ^39^, was also found beneficial to refine AlphaFold-Multimer models. Finally, AlphaFold-Multimer performs better than AlphaFold2 at protein-peptide complex prediction, and sampling a larger part of the conformational space by enforcing dropout at inference time in AlphaFold-Multimer further increased the quality of protein-peptide complex models ^40^.

In the present study, we investigate how best to use AlphaFold2 to make the leap from interaction networks to interfaces when dealing with binding partners containing intrinsically disordered regions (Figure 1a). We carefully develop an unbiased benchmark of 42 protein-peptide complexes sharing no similarity with any complex from the AlphaFold-Multimer training dataset and assess the performance of AlphaFold-Multimer on this dataset using different MSA schemes. We show that performance is limited when full-length protein sequences are used as input and considering delimited fragments increases the success rate. We set the fragment size at 200 amino acids in order to scan potential interacting regions within full-length sequences such as those derived from large-scale interactome data. Our study also raises the issue of prediction specificity, which may require the enumeration and ranking of potential anchoring sites, and assesses the usefulness of AlphaFold confidence scores in discriminating between possible binding regions. Finally, we show a synergistic effect of combining different MSA schemes and scores, allowing to reach 90% success rate on our benchmark dataset.

**Figure 1.**
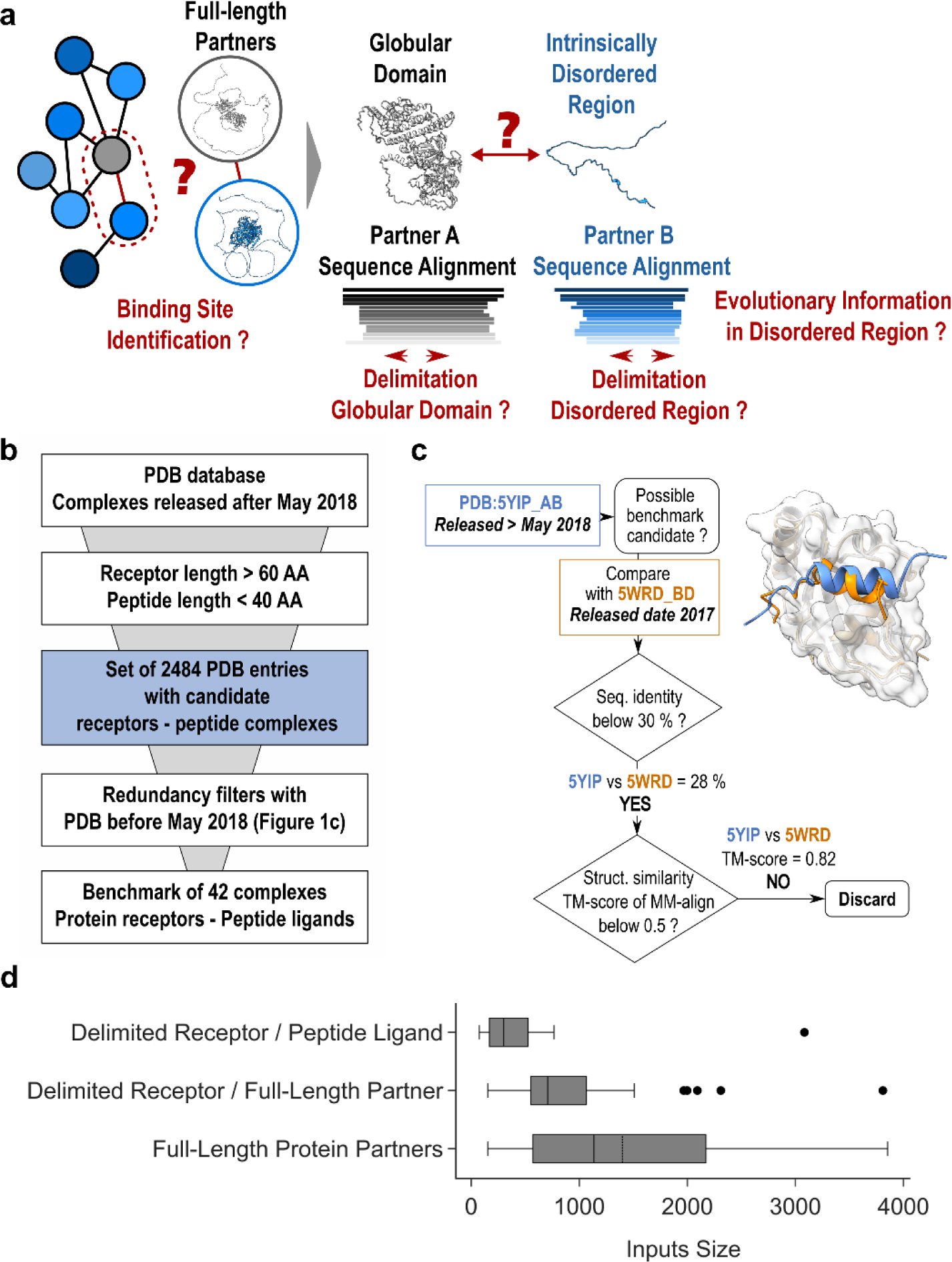
General presentation of the benchmark dataset. **(a)** Disentangling the complexity of a protein interaction network (sketched on the left) by analyzing binary interactions between a central grey protein and its blue binding partners can be complicated in case they contain intrinsically disordered regions **(b)** General pipeline to select the PDB entries that can be used as test complexes from those released after May 2018. They were required to share no sequence or structural redundancy with any of the complex structures that were used for AlphaFold2-Multimer training. **(c)** Example illustrating filters used to assess the lack of redundancy between the candidate complex and structures published before May 2018. Two filters were used, one based on sequence identity using a 30% seq. id. threshold and a second retrieving all complexes involving a receptor homolog using PPI3D ^50^ and checking for lack of interface structural similarity using MM-align ^52^. **(d)** Boxplots showing the cumulative size distribution of the 42 inputs (receptor+ligand) that were processed by AlphaFold2, either in protocols where sequences were delineated following the boundaries of the experimental structures or in those where full lengths of ligands and/or receptors were used.

## Results

### Selecting a test dataset of complexes not redundant with the training set of AF2-Multimer

To assess the performance of AlphaFold2 (AF2) in predicting the mode of association between a protein (hereafter called the receptor) and a small binding motif within a structurally disordered partner (the ligand), it is important to study cases of complexes that do not have homologs in the database on which AlphaFold2 has been trained. An example of how AF2 models may be biased by existing structures in the PDB is illustrated in Supp. Figure 1a. AF2-Multimer was trained on structures released until 30 Apr 2018. An analysis of the structures released after that date revealed that nearly 2,500 structures of complexes involving small protein motifs had been deposited in the PDB (Figure 1b). A large number of these structures have significant similarities in sequence or structure with structures released in the PDB before May 2018. Following a strict treatment of this sequence and structure redundancy (Figure 1c, see Methods), we isolated a set of 42 complexes involving a receptor and a small peptide ligand that could provide an unbiased estimate of AF2 performance in different conditions. Among the 42 complexes, we observed a diversity of subunit lengths (Figure 1d) and a representative occurrence of peptides with sizes ranging from 6 to 39 amino acids (Supp. Figure 1b) that are binding their receptors as helices, strands, coils or combination of those (Supp. Figure 1c).

AF2 relies on multiple sequence alignments whose evolutionary depth on the ligand peptide region may be limited due to the difficulty of identifying homologs from a short IDR sequence. Hence, for each of the proteins in this dataset, we used the full-length sequences of the protein partners to construct MSAs and subsequently delineate the interacting domains (Supp. Figure 2). These MSAs were combined to generate mixed co-alignments in which partner sequences belonging to the same species were paired while those with a single partner homolog present in a species were added as unpaired, similarly to ^41^. When the receptor and ligand are considered in their integrality, the overall length of the concatenated sequences is in majority between 1000 and 2000 amino acids, significantly larger than when the size of the inputs is delimited to the boundaries used for structural determination (Figure 1c). As a first analysis, we assessed whether AlphaFold2 was able to identify the correct binding site when proteins were considered in their full length. This is typical of a scenario where knowing that two proteins are interacting, we have no initial indication of which regions are involved.

### Success rates of AF2-Multimer for full-length and delimited input protein partners

For each run, 25 models were generated with AF2-Multimer parameters, following the reference protocol ^28^. The AF2 model confidence score (noted AF2 confidence score below), consisting of an 80:20 linear combination of ipTMscore and pTMscore, was used to rank the models and identify the best model. The accuracy of this best model was used to calculate the overall success rate for the 42 cases (see Methods). With full-length protein partners, we obtained a success rate of 40 % (Figure 2a), rather low with respect to that reported in the evaluation of AF2-Multimer for protein complexes, which was benchmarked using delimited sequence inputs ^28^. Analysis of the quality of the best model as a function of the size of the modeled assembly (Supp. Figure 3a) shows that the performance tends to decrease for large sizes above 1600 amino acids although it is still possible to observe good predictions above this size threshold. Below 1500 amino acids, the success rates do not appear correlated with the size of the assembly or the nature of the peptide secondary structure (Supp. Figure 3a).

**Figure 2.**
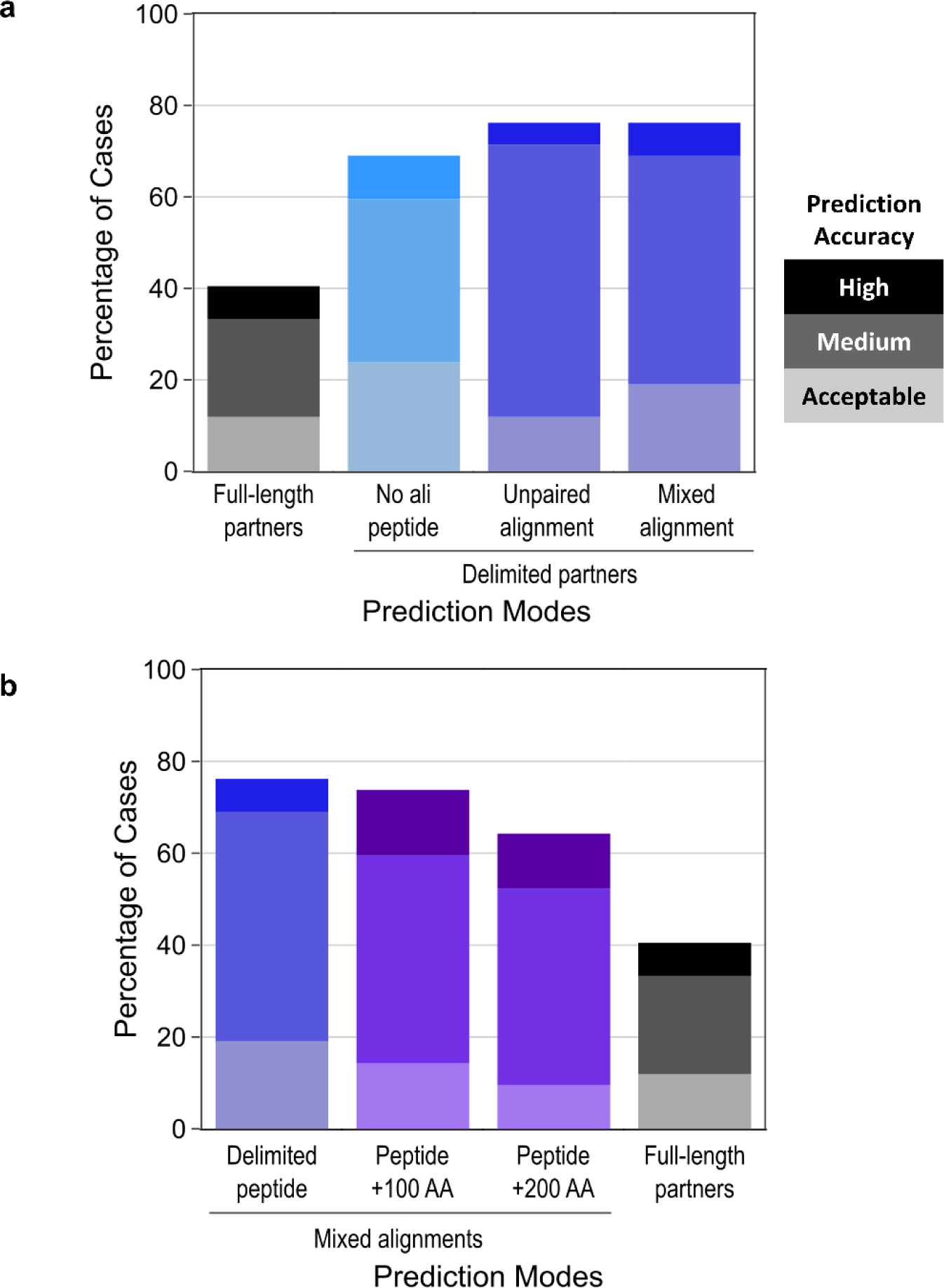
AlphaFold2-Multimer success rates on the benchmark dataset using different prediction modes. Stacked barplots reporting the success rates of AlphaFold2 prediction depending on the types of co-alignment used. All success rates are presented as the percentage of test cases in which the best AF2-Multimer model (best AF2 confidence score) is of Acceptable (light color), Medium (medium color) or High (dark color) quality according to the CAPRI criteria for protein-peptide complexes ^43^. **(a)** Success rates using (from left to right): full-length partners with a mixed alignment generation mode (grey grades), delimited partners with no evolutionary information for the peptide (cyan grades), delimited partners with unpaired co-alignment (blue grades), delimited partners with mixed alignment (blue grades). **(b)** Success rates using (from left to right): delimited partners (blue grades) (same as rightmost bar in panel a), peptides extended by 100 or 200 amino acids (purple grades), full-length partners (grey grades) (same as leftmost bar in panel a).

Next, the sequences of each binding partner were delimited according to their boundaries as observed in the experimental structure of the complex (Supp. Figure 2). When both the receptor and ligand were delimited, the overall success rate was much higher, reaching 76% of the 42 complexes correctly predicted (Figure 2a). In these first tests, the evolutionary information was integrated using the mixed co-alignment mode described above. We also tested alignment conditions in which the co-MSA is constructed from the same sequences but concatenated as an unpaired alignment. In this unpaired mode (Supp. Figure 2), the success rate remained similar at 76 % (Figure 2a), suggesting that homolog matching in the paired alignment does not provide a major gain. We assessed a third prediction mode in which no evolutionary information is added in the peptide region, as performed in ^37, 40^ (Supp. Figure 2). With this third approach, the performance obtained without evolutionary information on the peptide side remains high, with 69 % of correct models for the 42 cases (Figure 2a).

Such a good performance in the absence of any alignment associated with the peptide confirms that the properties of the binding site in the receptor are often sufficient to guide the interaction mode of the peptide ^37, 40^. Consistently, in a situation where the receptor is delimited but the ligand is considered in its full-length sequence, the performance drops back to a lower level of 50 %, even when using the MSA information on the ligand side (Supp. Figure 3b). Hence, one of the difficulties encountered by AF2 in dealing with large IDR-containing proteins lies in its ability to identify the correct interaction region within the partner protein.

The success rates calculated above were obtained by selecting only the model with highest AF2 confidence score among the 25 sampled models. Considering the entire set of 75 models (25 models for every complex in the three alignment conditions: mixed, unpaired, no_ali) highlights a significant Pearson’s correlation of 0.84 between the AF2 confidence score and the DockQ score, a commonly used metrics to rate the accuracy of modeled interfaces with respect to the reference complex ^42^ (Figure 3a). Grouping the models according to their CAPRI quality ranks (Acceptable/Medium/High) (Figure 3b) using the stringent criteria used for protein-peptide complexes ^43^ (see Methods) highlights that above an AF2 confidence score of 0.65, the predicted models are most often correct. There is also a minority of cases with a score between 0.4 and 0.65 that are found correct (in the Acceptable category) indicating that this twilight zone may be interesting to investigate if no alternative solution has been detected. In any case, the graphs on Figure 3a and 3b confirm that the AF2 confidence score (see Methods) can be used as a reliable proxy for estimating the reliability of a protein-peptide interaction prediction.

**Figure 3.**
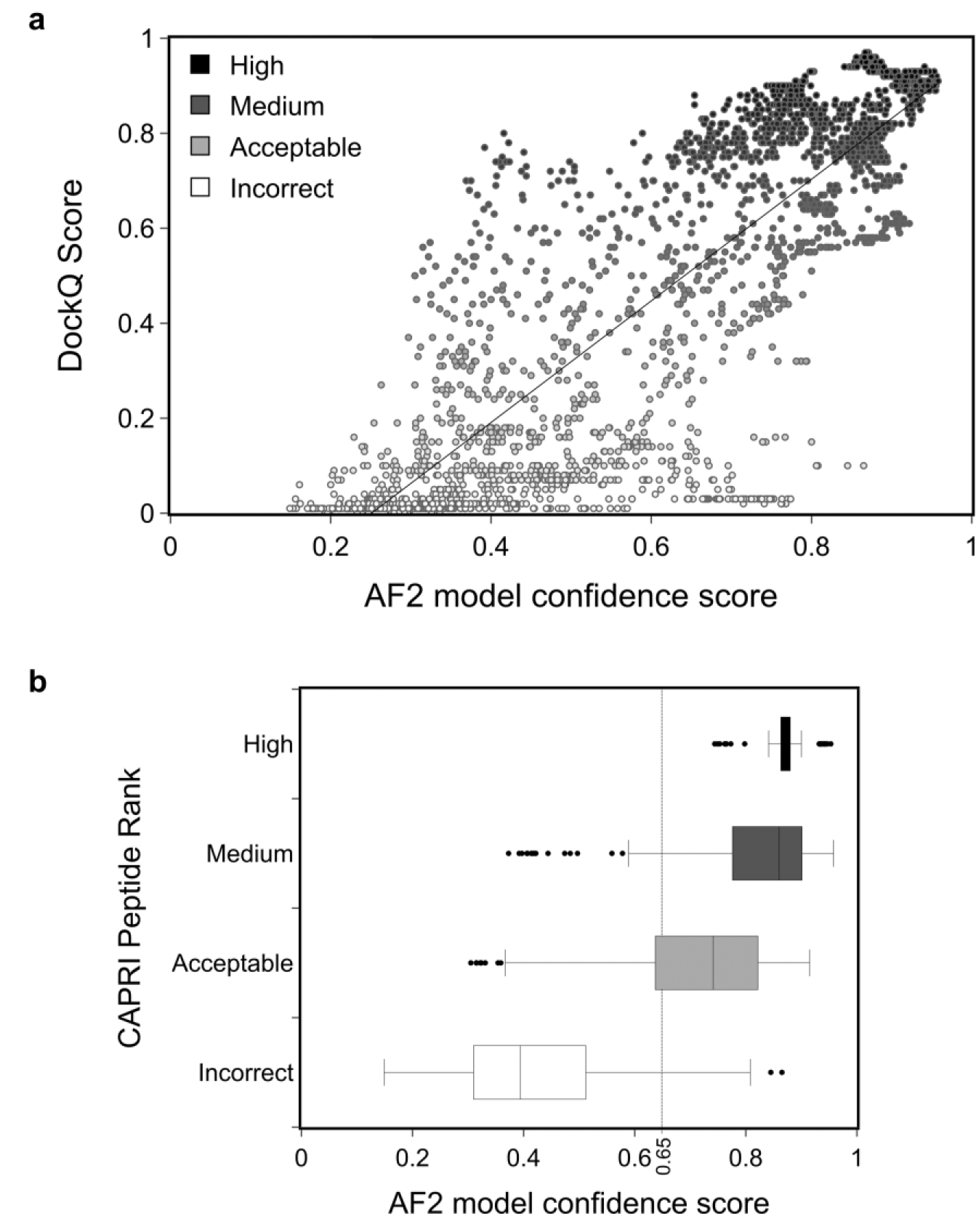
Model quality depending on the value of the AF2 model confidence score. (a) Distribution of DockQ scores ^42^ for 75 models for every binary protein-peptide complex (25 models in the three alignment conditions: mixed, unpaired, no_ali) as a function of the AF2 model confidence score. Data points are colored according to the model quality as rated by the DockQ score from low (white) to high (dark grey) values. Pearson’s correlation is 0.84. **(b)** Boxplots of the AF2 confidence score value distributions for the same set of models, split by model quality category according to the CAPRI protein-peptide criteria: High (dark grey), Medium (medium grey), Acceptable (light grey), Incorrect (white).

### Success rates of AF2-Multimer considering protein fragments of increasing size

When searching for an interaction site between two proteins, the region involved in the interaction is usually not known precisely. In order to use AF2 to carry out this task, and given the lower performance of AF2 with full-length proteins, we explored how AF2 predictions would be impacted by queries in which the bound motif is not perfectly delineated and is embedded in a larger fragment that may contain 100 or 200 additional amino acids. Extending the sequence containing the binding motif of each complex with up to 100 or 200 amino acids, and delimiting the alignments constructed in a mixed alignment mode (Supp. Figure 2), we obtained a decrease by 3 to 11 points with success rates of 73 % and 65 %, respectively for fragment size 100 and 200 (Figure 2b). The success rate of 65 %, obtained for cases where the fragment extends the peptide motif by 200 amino acids, is substantially higher than the 40 % obtained with full-length proteins. This result underscores the interest of fragment-based searching to identify potential interaction motifs between two partners and to predict their recognition mode. Previously (Figure 2a), we showed that the lack of evolution for the peptide was not very detrimental for a significant number of correct predictions (69 %). This trend is less pronounced when using fragments extended by 100 or 200 amino acids as shown in Supp. Figure 4. Without ligand alignment, there is a loss of performance of nearly 20 points, which highlights the importance of associating evolutionary information when the binding site identification involves a systematic search within larger fragments. For fragments of length 200, without evolutionary information for the ligand, the success rate is 45 %, almost as low as the success rates obtained for full-length proteins with evolution.

### Advantage of combining different alignment modes

The performance obtained using different alignment modes and input lengths suggests that some complexes can be correctly predicted regardless of the protocol used, while others may be sensitive to these input conditions. Overall, for 33 % (14 complexes), a correct model could be ranked first using the AF2 confidence score with any of the input conditions, even using full-length alignments (Supp. Figure 5). In contrast, other complexes could only be predicted correctly with a limited set of conditions (Figure 4a), suggesting a potential interest for combining different strategies. Instead of considering 25 models generated with every protocol, we analyzed a pool of 100 models generated with four different protocols and ranked them according to the highest AF2 confidence score. The resulting success rate improves significantly, rising up to 88 % (Figure 4b). The AF2 model confidence score is sufficiently correlated with the accuracy of the models that it can be used to identify correct assemblies in much larger model sets.

**Figure 4.**
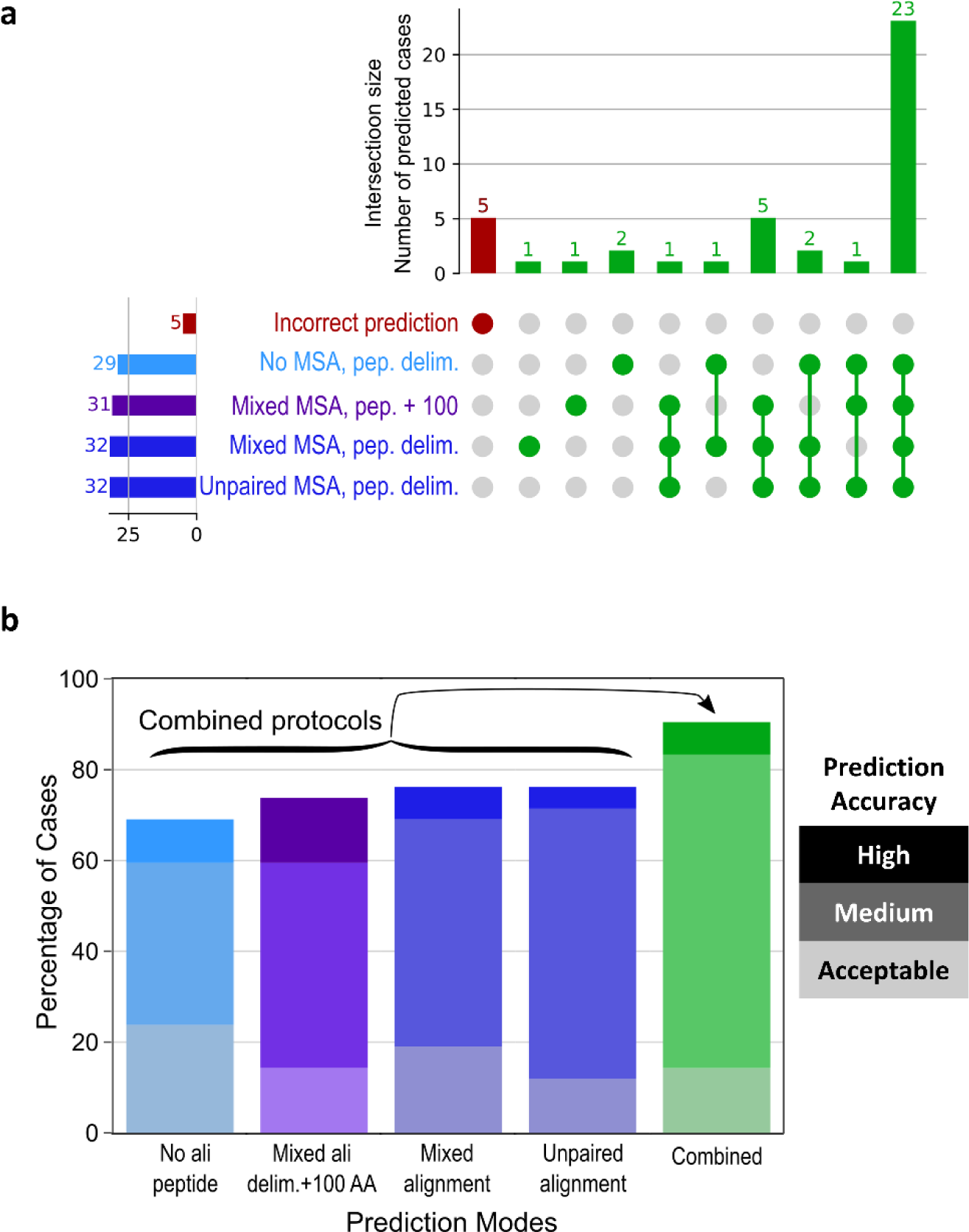
Complementarity of the predictions made in different prediction modes. (a) UpSet diagram ^58^ displaying the number of successful cases (out of 42) found by either a single or several prediction mode(s) among the following: delimited peptide with no peptide alignment, peptide extended by 100 residues with a mixed alignment, delimited peptide with a mixed alignment, delimited peptide with an unpaired alignment. 5 cases that can be identified in none of these conditions are highlighted in red. **(b)** Success rates for the four protocols shown in panel a (values are the same as in Fig 2) and for a combined protocol taking the best AF2 confidence score value out of 100 models (25 for each condition).

In an attempt to interpret the failures and successes of the tested protocols, we performed a detailed analysis of complexes that specifically succeeded with only a subset of the protocols and those that did not succeed with any. A typical case is when the conformation of the bound peptide is best predicted in the absence of evolutionary information. The absence of evolutionary information was found to be favorable for complexes such as 6ICV or 6YN0 that were not correctly predicted when MSA was added to the peptide. In the case of 6ICV (Figure 5a), the peptide (blue) is predicted to adopt a helix-and-turn conformation with high confidence when the evolutionary information of the MSA is included. This local structure is incompatible with the extended bound conformation. In contrast, in the absence of evolutionary information, the predicted structure of the peptide (light blue) is in very good agreement with the experimental structure (red), suggesting that the geometric constraints have been relaxed sufficiently for the peptide to sample an extended geometry that was well evaluated by the AF2 confidence score.

**Figure 5.**
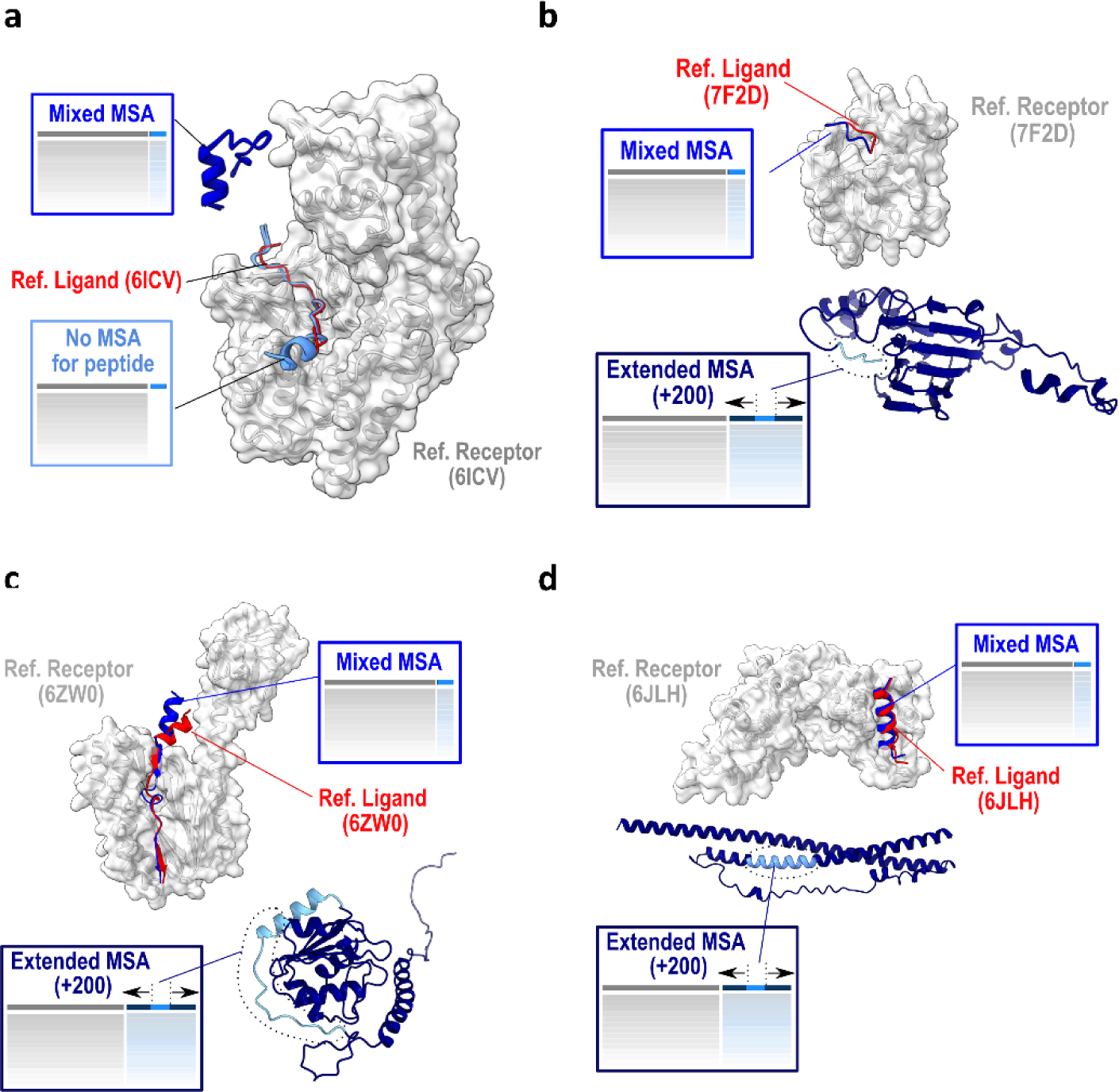
Detailed analysis of complexes that succeed with only a subset of the protocols. The receptor is represented as a grey surface, the native ligand as a red cartoon, the predicted peptides in shades of blue: bright blue for the predictions in mixed alignment mode, and light blue for the prediction with no peptide alignment (panel **a**) or for the peptide within the prediction of a fragment extended by 200 residues in dark blue (panels **b-d**). PDB identifiers of represented cases are 6ICV (**a**), 7F2D (**b**), 6ZW0 (**c**) and 6JLH (**d**).

Other differences between the tested protocols could be observed in case a motif, well predicted in a short fragment, was not correctly predicted in longer ones. This is observed in 5 cases including PDB cases 7F2D, 6ZW0 and 6JLH illustrated in Figure 5b, 5c and 5d, respectively. For these systems, considering the ligand peptide in the context of a larger fragment with 200 additional amino-acids (dark blue models) never led to a correct prediction by AF2, while the delimited peptides (blue) were always modeled in good agreement with the experimental reference structure (red). In almost all of these complexes, the origin of the failure in the larger fragments seems to be due to the presence of intramolecular contacts involving the peptide and surrounding regions. In the case of 7F2D and 6ZW0, the peptide is located in the vicinity of a globular domain with which it forms contacts of relatively low confidence. However, these appear to be sufficient to interfere with the generation of the native complex. In the third case, 6JLH, the binding peptide is embedded in a longer coiled-coil that masks the surface found to bind the receptor experimentally. This prediction would be consistent with the experimental study that showed the interaction to be observed only in specific physiological contexts ^44^. This example together with another case also involving long coiled-coils (7MU2) highlights the value of exploring different fragment lengths to reveal the appropriate binding epitopes. Therefore, in the cases where prediction performance varies between alignment content and delineation protocols, a common explanation is that the binding motif may be trapped or masked in a conformational state that prevents prediction of its correct binding mode.

Five cases out of 42 failed, regardless of the alignment protocol. In two of these cases (PDB: 6J08 and 7CZM), the receptor itself was not quite well folded, which may have made it difficult to sample a correct binding mode. For one case (PDB: 6A30) where none of the tested protocols converged to a correct model, we tested whether reducing the size of the receptor itself would help. We reran this case with the same alignments, testing if reduction in the size of the receptor could have an impact. Splitting the receptor as two inputs of similar size led to the generation of a correct model with a high AF2 confidence score with one of those inputs, reaching 0.8 when using the protocol with no peptide alignment but below 0.5 with all the other protocols. With this additional complex, the percentage of cases that could be predicted using AF2 rises above 90 %. Hence, there is room for further improvement by sampling simple alterations of the input MSAs and using the AF2 model confidence score as a guide for identification of the correct protocol.

### Specificity for similar binding motifs recognized by receptors

Out of the 42 cases in the test set, AF2 is able to correctly predict the binding mode of a peptide to its receptor without any evolutionary information for the peptide in 69 % of the cases. Such a performance suggests that the structural and evolutionary properties of the receptor match well with the peptide sequence irrespective of its conservation pattern. This calls into question the ability of AF2 to distinguish cognate binding peptides from non-binding ones. This issue may be particularly difficult in the challenging cases where two short fragments embedded in long disordered regions need to be discriminated while they tend to adopt a similar local conformation. To address this issue, we distinguished different classes of complexes based on the secondary structure adopted by the peptide in its bound conformation in order to create 83 challenging cross-partners predictions between 23 receptors and cognate or non-cognate ligands selected from the 42 cases of our test set. We then assessed whether AF2 could specifically predict the binding mode of receptors with their respective peptides and distinguish them from potentially misleading peptides taken from unrelated structures but sharing similar bound conformations.

In total, 7 categories of peptide conformations were considered (Supp. Figure 6). We distinguished those binding through a small, medium, or long helix, those showing no canonical secondary structure and those binding through the formation of a combination of helix and strand or a single or two beta-strands (Supp. Figure 6a to 6g). To run the cross-partners interaction tests, we used the protocol with no MSA in the peptide region. Over the 23 selected cases for cross-partners analysis, 16 were successfully predicted by AF2 (70 %) in agreement with the performance obtained with this protocol on the 42 test cases. In Figure 6a, the distribution of AF2 confidence scores obtained for the models rated as correct using the CAPRI protein-peptide criteria (darker blue distribution) differs significantly from the distribution of the scores obtained with non-native peptide ligands (light blue distribution). The AF2 confidence score of the specific peptide was superior to any of the non-specific peptides in 15 out of the 16 correctly predicted complexes. Figure 6b illustrates one of these 15 cases, using the receptor of 7CFC, highlighting that even if the non-specific peptides tend to interact in the same region as the specific one, the AF2 confidence score is higher for the specific peptide (reaching 0.75) and can be used as a proxy to discriminate between several likely binders.

**Figure 6.**
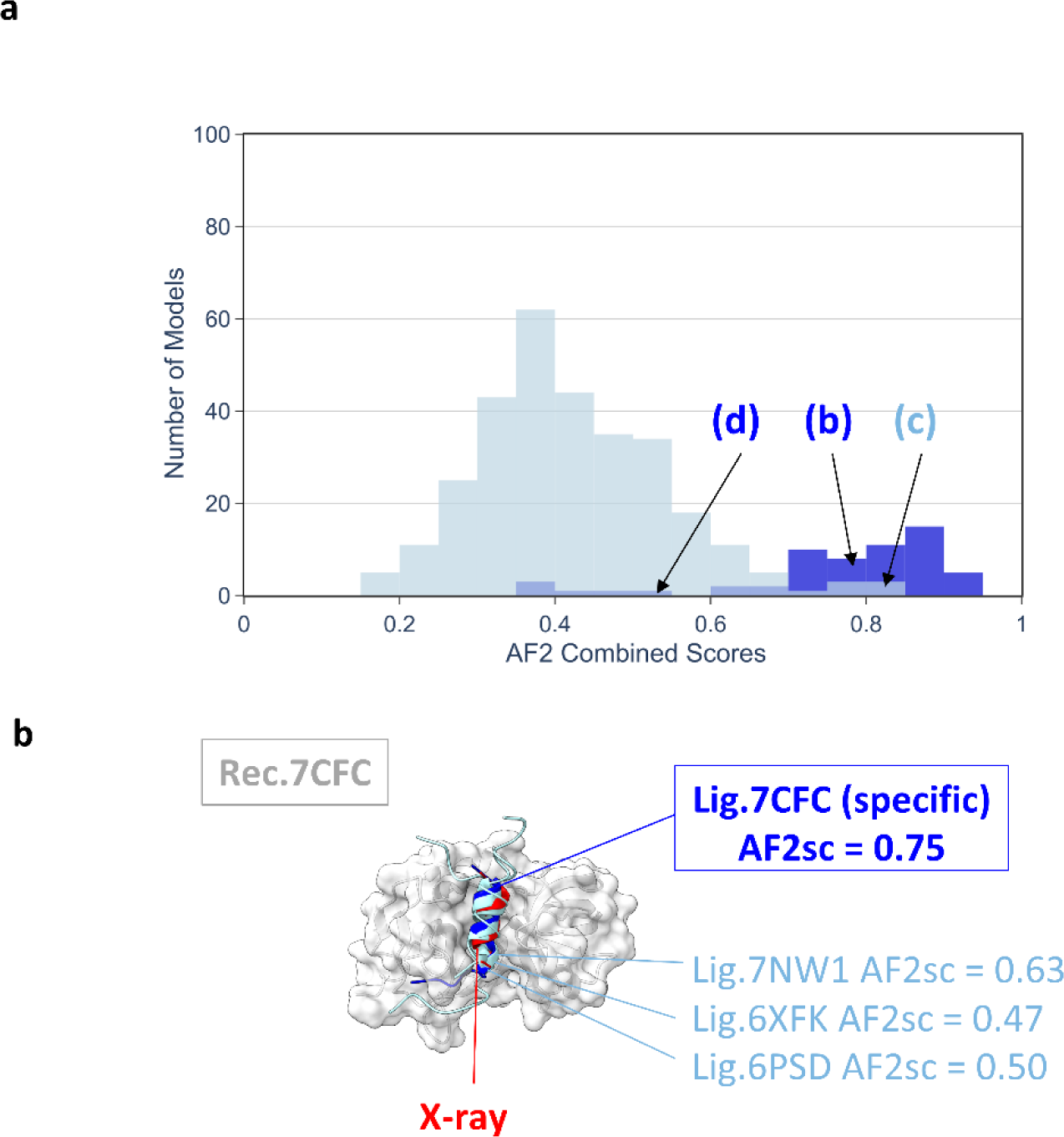
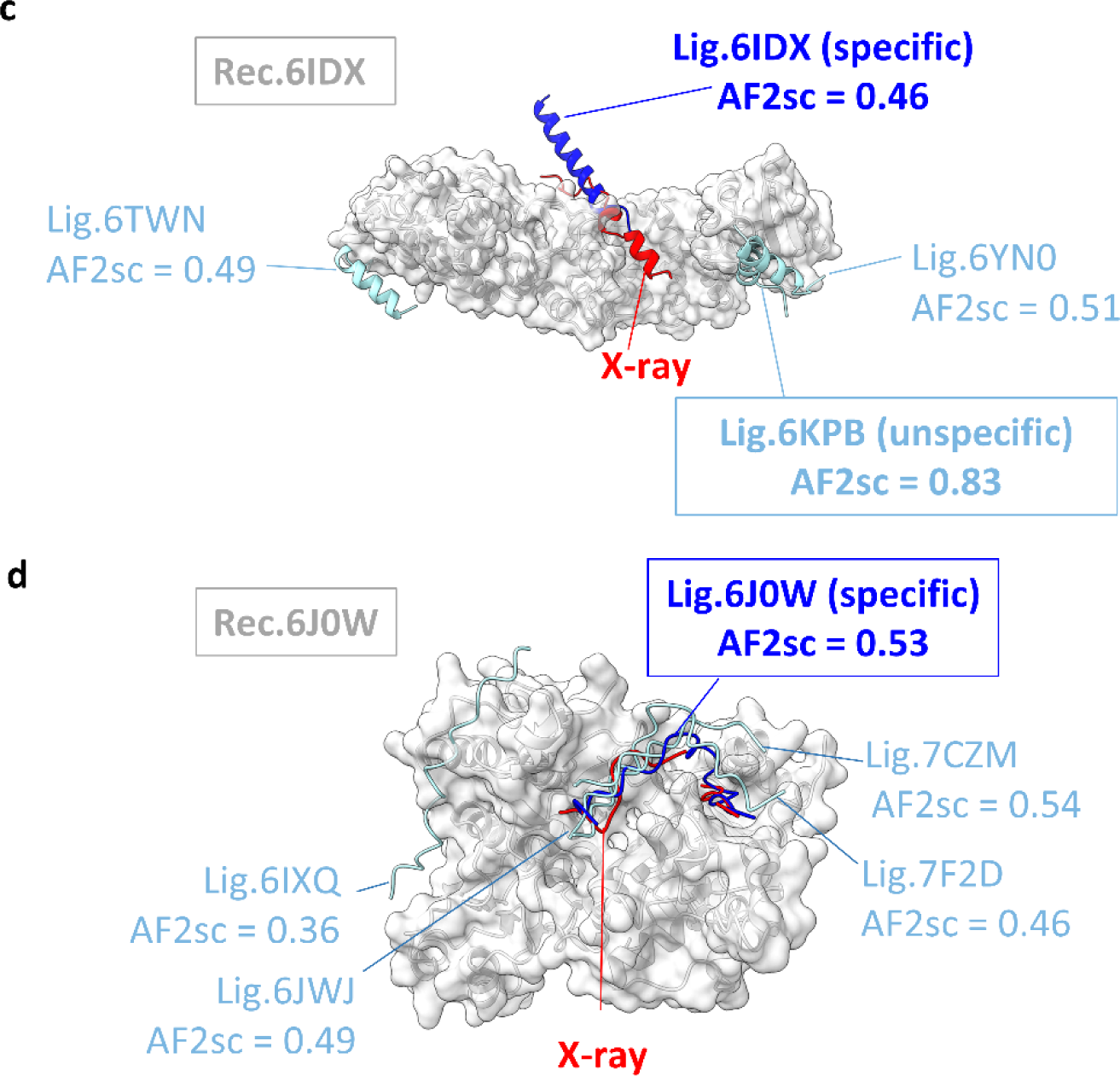
Cross-partners evaluation assessing the specificity of the binding predictions. (a) Distribution of AF2 confidence scores obtained for the models involving the native peptide and rated as correct using the CAPRI protein-peptide criteria (darker blue distribution) for 16 out of 23 cases selected for cross-partners evaluation and for the models obtained with non-native peptide ligands (light blue distribution). Cross-partners predictions were performed using delimited partners with no evolutionary information for the peptide. Specific predictions illustrated in panels **(b-d)** are drawn from the relevant distributions at the indicated score values. (**b-d**) Illustration of specific cases discussed in the text for native PDB identifiers: 7CFC (**b**), 6IDX (**c**), 6J0W (**d**). The receptor is shown as a grey surface, the native peptide as a red cartoon, the best predicted model involving the native peptide in bright blue cartoon and the best predicted models involving non-native peptides in light blue. AF2sc values indicate the best AF2 confidence score values obtained for each complex.

Based on the distribution in Figure 6a, a minority of models (approximately 10%) would give a misleading assignment for AF2 confidence scores greater than 0.6 and could prevent identification of a correct binding site. This is illustrated by the case of the 6IDX complex in Figure 6c in which an incorrect binder (6KPB Ligand) is predicted to form a complex with the receptor with high confidence (as indicated by an AF2 confidence score of 0.83) while the specific ligand was not correctly predicted (AF2 confidence score=0.46 and wrong binding mode).

Last, there are also alternative situations as illustrated in Figure 6d in which the score of the specific binder is mild (below 0.6) but still among the highest scores obtained in the set of potential binders. This was observed for 3 of the 16 cases where the specific receptor-ligand pair was correctly predicted by AF2 (6YN0 in Supp. Figure 6c and 7F2D, 6J0W in Supp. Figure 6f). With the 6J0W receptor (Figure 6d), the AF2 confidence score of the specific ligand is 0.53, whereas it reaches 0.54 with another non- specific ligand of 7CZM. Such a situation highlights the specificity issue that may arise in the case where the peptide is not accurately modeled in the receptor binding site. It can be noted in Supp. Figure 6f that the misleading 7CZM ligand tends to have higher AF2 confidence scores than the other ligands on average with all the non-specific receptors. Such promiscuity indicates the risk that some sequences may systematically bias the specificity analysis and that normalization or the use of an alternative scoring scheme might be useful to further disentangle the complexity of protein-protein interaction networks involving unstructured regions.

## Discussion

AF2 has shown remarkable performance for predicting the structure of multi-subunit machineries only known so far through PPI maps ^26, 27, 30^. In this study, we explored the potential of AlphaFold2 to further exploit the wealth of data contained in proteomics experiments and to enable a more comprehensive characterization of protein-protein interaction networks. We focused on interactions mediated by unstructured regions that are a cornerstone of the functional and dynamic organization of most cellular processes. The capacity of AF2 to perform well with small disordered regions binding a structured domain was established on different datasets ^18, 37, 40^ built from the structures available in the PDB ^45^. However, in most proteomics experiments, the boundaries of the interacting regions are not precisely known. In addition, physical interactions can be indirect and it is crucial to disentangle the regions involved in direct interactions.

To further assess the use of AF2 for this purpose, we built a dataset of complexes consisting of a folded receptor bound to a short protein fragment and evaluated several protocols representative of challenges faced following proteomics analyses. Because AlphaFold2-multimer was trained on complexes whose structures were published before May 2018, we carefully removed any homologs of the complex to ensure that our conclusions could not be biased by similarities in sequence or 3D geometry with the training dataset. For the 42 test cases selected in the benchmark, we first evaluated the ability of AF2 to discriminate the binding site when proteins are provided in their full length as in the output of a proteomic experiment. We achieved a success rate of 40% and noted that above 1600 amino acids, the method gave poorer predictions, with the exception of two impressive cases above 2500 amino acids. The use of input fragments delimited as in the experimental structures significantly increased performance by more than 35 points and the combination of different MSA construction modes led to an overall success rate of 90%. If the binding region is unknown, scanning multiple small peptides can be computationally demanding and we found that a reasonable trade-off in accuracy could be achieved with a fragment length of about 100 amino acids. For those lengths, evolutionary information was key to reaching the best results. For larger fragments involving more than 200 amino- acids, a decrease in performance was observed, not necessarily due to an input size greater than 1600 amino acids. Our analysis rather suggests that the drop may also originate from intramolecular contacts that tend to mask the binding region or hinder the sampling of the bound conformation.

In the case of delimited peptides, it is remarkable that evolutionary information in the peptide region did not prove to be as crucial as for longer fragments for generating accurate models and scoring them reliably. We found that in specific cases where the bound conformation of the peptide was rather extended, the addition of evolutionary information was even detrimental to the identification of a correct solution. Such a detrimental effect of MSA was also reported in ^46^ for the structural prediction of complexes between MHC receptors and various sets of short peptides by AF2. In these systems, the local conformation of the bound peptides is also fully extended. Our analysis suggests that the inclusion of the MSA for the disordered short peptide may lock the local conformations of the peptides and prevent them from adopting a different bound conformation. However, we also found that as the size of the disordered fragment increases beyond 100 amino acids, the requirement for evolutionary information becomes more critical. In any case, sampling these different possibilities was considered worthwhile, as the AF2 confidence score is sufficiently reliable to pick out the correct solution among those sampled.

Beyond the remarkable ability of AF2 to generate correct conformations of protein-peptide complexes, we confirmed the reliability of the combined ipTMscore and pTMscore as an estimate of model accuracy. We also evaluated AF2 as a tool to discriminate a native ligand from other ligands potentially difficult to discriminate because adopting the same local conformation. The obtained results were satisfactory in a majority of cases where the AF2 confidence score correctly singled out the native binding peptide, but also highlighted several misleading situations that call for vigilance in the exploitation of specificity results. It certainly should be possible to reinforce the applicability of AF2 for the exploitation of more complex interactomes in which the interaction with unstructured regions plays a major role. Recent efforts in that direction have shown that AF2 parameters which were trained only with positive examples could be further fine-tuned for specificity combining positive and negative examples of receptor-peptide interactions ^46^. So far, this fine-tuning was achieved in a receptor-specific manner focusing on MHC, PDZ or SH3 domains, but it might be expanded further to address other specificity issues.

The ability of AF2 to discriminate the native peptide from similar alternative binders when the native bound conformation is correctly predicted supports the conclusions that an energetic function of the protein structure has been learned by AF2 independently of evolutionary information ^47^. This ability to discriminate specific native binders is also consistent with the principle of using AF2 to design high- affinity binders for their targets ^48, 49^. The fact that with larger fragments (>200), the ability to identify the correct binding site decreases significantly and requires evolutionary information is also in agreement with the proposal that AF2 needs coevolution data to search for global minima in the learned function ^47^. To progress from interactomes to the identification of all potential binding sites within disordered regions, a robust strategy would benefit from systematically scanning fragments of sequences of limited length and sampling different types of evolutionary information, such as the four combined in this study.

## Methods

### Building the dataset of protein-peptide complexes non-redundant with the AF2 training structural dataset

An initial list of protein-peptide complexes was retrieved from the PDB server ^45^ on April 1, 2022 with the following request: 1) Release date after May 1st, 2018 to exclude complexes present in the AlphaFold2 training set; 2) The longest protein (called the receptor) must contain at least 60 amino acids and the smallest chain (called the peptide) must contain at most 40 amino acids. 3) The ‘Number of Polymer Instances (Chains) per Assembly’ has to be between 2 and 4 and should contain heteromeric assemblies. 4) The assemblies should not contain RNA or DNA chains. The initial request led to 2484 potential candidates. Using a sequence identity threshold of 30 %, we discarded all candidates for which a homolog of the receptor protein was released before May 1st, 2018 and bound to a ligand partner in the same region. From the list of selected candidates, an additional filter was used to check the absence of redundant assembly modes. For each of the selected complexes, the receptor sequence was used as a query of the PPI3D server ^50^, in single sequence mode, to recover all the PDB codes of complexes involving homologs of the receptor (date of PPI3D query August 2022, on the PDB updated July 20, 2022). In PPI3D, distant receptor homologs were retrieved using PSI-BLAST ^51^ with 2 iterations and an E-value cutoff of 0.002. For every candidate complex, PPI3D provided a detailed list of PDB codes with the chain ids involving the receptor or its homolog. We used the full list of interactions provided by PPI3D, except when it exceeded 2500 interfaces in which case the clustered list was chosen (95% sequence similarity and 50% similarity for residues in the binding region). Only the interfaces annotated as ‘hetero’ or ‘hetero-peptide’ released before May 1, 2018 were considered as potentially redundant. Their structures were compared to the candidate complex using the MM-align program (Version 20191021) ^52^ (option “-a”) and the maximum of the three TM-scores calculated was considered. Receptor-peptide candidates with a TM-score greater than 0.5 with any other potentially redundant interface extracted from the PPI3D results were considered redundant with a previously known structure and were discarded. This latter condition only applied to structures for which MM-align successfully aligned at least 5 consecutive amino acids on the ligand side (detected by ‘:’ in the output pair alignment corresponding to residue distance pairs < 5.0 Angstrom), otherwise the interface was not considered redundant. In the end, we retained a set of 42 receptor-peptide cases to form the reference database.

### Generation of the alignments for the 42 database cases

Sequences of all the chains in the dataset of 42 complexes were retrieved from the UniProt database ^53^ and were submitted to three iterations of MMseqs2 ^54^ against the uniref30_2103 database ^41^. The resulting alignments were filtered using hhfilter ^55^ using parameters (‘id’=100, ‘qid’=25, ‘cov’=50) and the taxonomy assigned to every sequence keeping only one sequence per species. Full-length sequences in the alignments were then retrieved and the sequences were realigned using MAFFT ^56^ with the default FFT-NS-2 protocol. To build the so-called mixed co-alignments, sequences in the alignment of individual partners were paired according to their assigned species and left unpaired in case no common species were found ^41^. Unpaired alignments were obtained by unpairing the mixed alignments and alignments with no evolutionary information for the ligand were obtained by leaving the ligand region as a single sequence. The concatenated multiple sequence alignments (MSA) were delimited using as input the MSA generated with the three protocols described above (mixed, unpaired, single sequence). The sequence delimitations as defined in the SEQRES PDB parameter were used to delineate the receptor and the peptide. In case the receptor was a heteromer or assembled as a homodimer, the full complex was modeled. To generate the models extended by 100 or 200 amino acids, the peptide sequence was extended in both directions unless a chain end was encountered in which case the extension was pursued in only one direction.

### Generation of input data for cross-partners evaluation

To generate the dataset mixing receptors and their non-cognate ligands, a subset of complexes that could be clustered according to the similarity of the type and length of the secondary structure of their ligand (reported in Supp. Table 1) was defined (Supp. Table 2). We selected 23 complexes with a monomeric receptor and a ligand that could be clustered into one of the 7 groups distinguished in Supp Table 2. The MSA of each receptor was concatenated with each ligand in the same cluster without adding MSA information on the ligand side. These alignments were used as input to generate structural models by AlphaFold2 following the protocol described below.

### Generation of the structural models

The concatenated MSAs were used as input to run 5 independent runs of the AlphaFold2 algorithm with 3 recycles each ^15, 28^ generating 5 structural models using a local version of ColabFold v1.3 ^41^ with the Multimer v.2.2 model parameters ^28^ on NVidia A100 GPUs. Four scores were provided by AlphaFold2 to rate the quality of the models, the pLDDT, the pTMscore ^15^, the ipTMscore and the model confidence score (weighted combination of pTM and ipTM scores with a 20:80 ratio) ^28^. The scores obtained for all the generated models are reported in Supp. Table 3.

### Evaluation and visualization of the structural models

The structural models generated with every alignment protocols were compared to their reference structure defined in Supp. Table 1. The models were first delimited as in the reference experimental structure to ensure proper superposition of receptors and ligands. The accuracy of the models was assessed using two related measures (i) the DockQ score, which provides a continuous value between 0 and 1, with limits of 0.23, 0.49, and 0.8 defining Acceptable, Medium, and High quality thresholds for protein-protein complexes ^42^ (ii) the more stringent conditions established by the CAPRI community to rate the specific cases of receptor-peptide complexes using ligand and interface Root-Mean-Square Deviation (L- and iRMSD) and the Fraction of native contacts (fnat). Ranks are assigned depending on the following criteria: High (fnat in [0.8, 1.0] and (L-RMSD ≤ 1.0 Å or iRMSD ≤ 0.5 Å)), Medium (fnat in [0.5, 0.8] and (L-RMSD ≤ 2.0 Å or iRMSD ≤ 1.0 Å) or fnat in [0.8, 1.0] and (L-RMSD>1.0 Å and iRMSD>0.5 Å)) and Acceptable (fnat in [0.2, 0.5] and (L-RMSD ≤ 4.0 Å or iRMSD ≤ 2.0 Å) or fnat [0.5, 1.0] and (L- RMSD>2.0 Å AND iRMSD>1.0 Å)) ^43^. Additional analyses were performed following the standard metrics calculated by CAPRI assessors to rate the similarity between the models and their reference structure (such as the fraction of interface residues FRIR or the fraction of non-native contacts FRNNAT) and are also available in Supp. Table 3. 3D structures were visualized and represented using ChimeraX ^57^.

## Data availability

The reference PDB files of the 42 test cases, the multiple sequence alignments built for all ten protocols and the corresponding PDB files of the predicted models are available from ZENODO under accession DOI code 10.5281/zenodo.7838024.

## Code availability

The code for processing, analyzing and visualizing the results is available at: https://github.com/i2bc/SCAN_IDR

## Author contributions

HB, JA and RG designed the study. HB built the dataset and developed the scripts to generate the alignments, ran AlphaFold2 and analyze the outputs. JA and RG jointly supervised the work and wrote the manuscript together with HB.

## Acknowledgements

The work was supported by grants from Agence Nationale de la Recherche (ANR-21-CE44-0009-01 and ANR-18-CE45-0005-01 ESPRINet).

## References

1. Van Roey K, et al. Short linear motifs: ubiquitous and functionally diverse protein interaction modules directing cell regulation. Chem Rev 114, 6733–6778 (2014).

2. Uyar B, Weatheritt RJ, Dinkel H, Davey NE, Gibson TJ. Proteome-wide analysis of human disease mutations in short linear motifs: neglected players in cancer? Mol Biosyst 10, 2626–2642 (2014).

3. Uversky VN. Intrinsic Disorder, Protein-Protein Interactions, and Disease. Adv Protein Chem Struct Biol 110, 85–121 (2018).

4. Kumar M, et al. The Eukaryotic Linear Motif resource: 2022 release. Nucleic Acids Res 50, D497–D508 (2022).

5. Jehl P, Manguy J, Shields DC, Higgins DG, Davey NE. ProViz-a web-based visualization tool to investigate the functional and evolutionary features of protein sequences. Nucleic Acids Res 44, W11–15 (2016).

6. Erdos G, Pajkos M, Dosztanyi Z. IUPred3: prediction of protein disorder enhanced with unambiguous experimental annotation and visualization of evolutionary conservation. Nucleic Acids Res 49, W297–W303 (2021).

7. Cafarelli TM, Desbuleux A, Wang Y, Choi SG, De Ridder D, Vidal M. Mapping, modeling, and characterization of protein-protein interactions on a proteomic scale. Curr Opin Struct Biol 44, 201–210 (2017).

8. Elhabashy H, Merino F, Alva V, Kohlbacher O, Lupas AN. Exploring protein-protein interactions at the proteome level. Structure 30, 462–475 (2022).

9. Ghadie MA, Coulombe-Huntington J, Xia Y. Interactome evolution: insights from genome-wide analyses of protein-protein interactions. Curr Opin Struct Biol 50, 42–48 (2018).

10. Wan C, et al. Panorama of ancient metazoan macromolecular complexes. Nature 525, 339–344 (2015).

11. Holguin-Cruz JA, Foster LJ, Gsponer J. Where protein structure and cell diversity meet. Trends Cell Biol, (2022).

12. Mosca R, Pache RA, Aloy P. The role of structural disorder in the rewiring of protein interactions through evolution. Mol Cell Proteomics 11, M111 014969 (2012).

13. Andreani J, Quignot C, Guerois R. Structural prediction of protein interactions and docking using conservation and coevolution. WIREs Computational Molecular Science 10, e1470 (2020).

14. Gibson TJ, Dinkel H, Van Roey K, Diella F. Experimental detection of short regulatory motifs in eukaryotic proteins: tips for good practice as well as for bad. Cell Commun Signal 13, 42 (2015).

15. Jumper J, et al. Highly accurate protein structure prediction with AlphaFold. Nature 596, 583–589 (2021).

16. Tunyasuvunakool K, et al. Highly accurate protein structure prediction for the human proteome. Nature 596, 590–596 (2021).

17. Ruff KM, Pappu RV. AlphaFold and Implications for Intrinsically Disordered Proteins. J Mol Biol 433, 167208 (2021).

18. Akdel M, et al. A structural biology community assessment of AlphaFold2 applications. Nat Struct Mol Biol 29, 1056–1067 (2022).

19. Wilson CJ, Choy WY, Karttunen M. AlphaFold2: A Role for Disordered Protein/Region Prediction? Int J Mol Sci 23, (2022).

20. Seoane B, Carbone A. Soft disorder modulates the assembly path of protein complexes. PLoS Comput Biol 18, e1010713 (2022).

21. Bryant P, Pozzati G, Elofsson A. Improved prediction of protein-protein interactions using AlphaFold2. Nat Commun 13, 1265 (2022).

22. Yin R, Feng BY, Varshney A, Pierce BG. Benchmarking AlphaFold for protein complex modeling reveals accuracy determinants. Protein Sci 31, e4379 (2022).

23. Si Y, Yan C. Protein complex structure prediction powered by multiple sequence alignments of interologs from multiple taxonomic ranks and AlphaFold2. Brief Bioinform 23, (2022).

24. Ghani U, et al. Improved Docking of Protein Models by a Combination of Alphafold2 and ClusPro. bioRxiv, 2021.2009.2007.459290 (2022).

25. Gao M, Nakajima An D, Parks JM, Skolnick J. AF2Complex predicts direct physical interactions in multimeric proteins with deep learning. Nat Commun 13, 1744 (2022).

26. Burke DF, et al. Towards a structurally resolved human protein interaction network. Nat Struct Mol Biol, (2023).

27. O’Reilly FJ, et al. Protein complexes in cells by AI-assisted structural proteomics. Mol Syst Biol 19, e11544 (2023).

28. Evans R, et al. Protein complex prediction with AlphaFold-Multimer. bioRxiv, 2021.2010.2004.463034 (2022).

29. Baek M, et al. Accurate prediction of protein structures and interactions using a three-track neural network. Science 373, 871–876 (2021).

30. Humphreys IR, et al. Computed structures of core eukaryotic protein complexes. Science 374, eabm4805 (2021).

31. Del Alamo D, Sala D, McHaourab HS, Meiler J. Sampling alternative conformational states of transporters and receptors with AlphaFold2. Elife 11, (2022).

32. Stein RA, McHaourab HS. SPEACH_AF: Sampling protein ensembles and conformational heterogeneity with Alphafold2. PLoS Comput Biol 18, e1010483 (2022).

33. Yu DQ, Chojnowski G, Rosenthal M, Kosinski J. AlphaPulldown-a python package for protein-protein interaction screens using AlphaFold-Multimer. Bioinformatics, 2022.2008.2005.502961 (2022).

34. Iserte JA, Lazar T, Tosatto SCE, Tompa P, Marino-Buslje C. Chasing coevolutionary signals in intrinsically disordered proteins complexes. Sci Rep 10, 17962 (2020).

35. Ciemny M, et al. Protein-peptide docking: opportunities and challenges. Drug Discov Today 23, 1530–1537 (2018).

36. Schueler-Furman O, London N. Modeling Peptide-Protein Interactions. Methods and Protocols. Humana Press (2017).

37. Tsaban T, Varga JK, Avraham O, Ben-Aharon Z, Khramushin A, Schueler-Furman O. Harnessing protein folding neural networks for peptide-protein docking. Nat Commun 13, 176 (2022).

38. Johansson-Akhe I, Wallner B. InterPepScore: A Deep Learning Score for Improving the FlexPepDock Refinement Protocol. Bioinformatics, (2022).

39. Alam N, Goldstein O, Xia B, Porter KA, Kozakov D, Schueler-Furman O. High-resolution global peptide-protein docking using fragments-based PIPER-FlexPepDock. PLoS Comput Biol 13, e1005905 (2017).

40. Johansson-Akhe I, Wallner B. Improving peptide-protein docking with AlphaFold-Multimer using forced sampling. Front Bioinform 2, 959160 (2022).

41. Mirdita M, Schutze K, Moriwaki Y, Heo L, Ovchinnikov S, Steinegger M. ColabFold: making protein folding accessible to all. Nat Methods 19, 679–682 (2022).

42. Basu S, Wallner B. DockQ: A Quality Measure for Protein-Protein Docking Models. PLoS One 11, e0161879 (2016).

43. Lensink MF, Velankar S, Wodak SJ. Modeling protein-protein and protein-peptide complexes: CAPRI 6th edition. Proteins 85, 359–377 (2017).

44. Qin J, et al. Structural and mechanistic insights into secretagogin-mediated exocytosis. Proc Natl Acad Sci U S A 117, 6559–6570 (2020).

45. Burley SK, et al. RCSB Protein Data Bank (RCSB.org): delivery of experimentally-determined PDB structures alongside one million computed structure models of proteins from artificial intelligence/machine learning. Nucleic Acids Res 51, D488–D508 (2023).

46. Motmaen A, Dauparas J, Baek M, Abedi MH, Baker D, Bradley P. Peptide-binding specificity prediction using fine-tuned protein structure prediction networks. Proc Natl Acad Sci U S A 120, e2216697120 (2023).

47. Roney JP, Ovchinnikov S. State-of-the-Art Estimation of Protein Model Accuracy Using AlphaFold. Phys Rev Lett 129, 238101 (2022).

48. Chang L, Perez A. Ranking Peptide Binders by Affinity with AlphaFold. Angew Chem Int Ed Engl, e202213362 (2022).

49. Bryant P, Elofsson A. EvoBind: in silico directed evolution of peptide binders with AlphaFold. bioRxiv, 2022.2007.2023.501214 (2022).

50. Dapkunas J, Timinskas A, Olechnovic K, Margelevicius M, Diciunas R, Venclovas C. The PPI3D web server for searching, analyzing and modeling protein-protein interactions in the context of 3D structures. Bioinformatics 33, 935–937 (2017).

51. Altschul SF, et al. Gapped BLAST and PSI-BLAST: a new generation of protein database search programs. Nucleic Acids Res 25, 3389–3402 (1997).

52. Mukherjee S, Zhang Y. MM-align: a quick algorithm for aligning multiple-chain protein complex structures using iterative dynamic programming. Nucleic Acids Res 37, e83 (2009).

53. UniProt C. UniProt: the Universal Protein Knowledgebase in 2023. Nucleic Acids Res 51, D523–D531 (2023).

54. Steinegger M, Soding J. MMseqs2 enables sensitive protein sequence searching for the analysis of massive data sets. Nat Biotechnol 35, 1026-1028 (2017).

55. Steinegger M, Meier M, Mirdita M, Vohringer H, Haunsberger SJ, Soding J. HH-suite3 for fast remote homology detection and deep protein annotation. BMC Bioinformatics 20, 473 (2019).

56. Katoh K, Standley DM. MAFFT multiple sequence alignment software version 7: improvements in performance and usability. Mol Biol Evol 30, 772–780 (2013).

57. Pettersen EF, et al. UCSF ChimeraX: Structure visualization for researchers, educators, and developers. Protein Sci 30, 70–82 (2021).

58. Lex A, Gehlenborg N, Strobelt H, Vuillemot R, Pfister H. UpSet: Visualization of Intersecting Sets. IEEE Transactions on Visualization and Computer Graphics 20, 1983–1992 (2014).

