## Supplementary material for "From interaction networks to interfaces: Scanning intrinsically disordered regions using AlphaFold2": Supp Tables and Figures

### Supplementary Table Legends

**Supplementary Table 1. Table of the 42 test cases with their main characteristics**

**Supplementary Table 2. Table of the 7 clusters that were used to sample the cross-partner interactions**

**Supplementary Table 3. Table of all the AF2 and structural evaluation scores for all the models generated in the study of the 42 test cases with 10 different alignment protocols (provided as excel file)**

### Supplementary Figures Legends

**Supplementary Figure 1. Benchmark dataset construction.**

**(a)** (Left) Structure of the intrinsically disordered region (IDR) of human MCM2 (orange) with boundaries [68-125] as experimentally resolved in complex with histones H3 (blue) and H4 (cyan) (PDB: 5BNV) compared (right) to the full AlphaFold model of MCM2 extracted from the AlphaFold database and colored orange in the same boundaries [68-125]. No other structure of this region is available in the PDB. The similarity between the local structures in this stretch highlights that the conformations in the model may be strongly inspired from experimental structures used in the training process of AlphaFold. **(b)** Boxplots representing the size distribution of the 42 peptides in the benchmark data set. **(c)** Classification of the different types of secondary structures observed in the ligands of the 42 complexes in the benchmark database. Four categories were distinguished, peptide structures comprising only helical secondary structure (Helix) or only strand(s) (Strand), peptides binding in the absence of a canonical secondary structure (Coil), and those adopting more complex combinations of the three elements mentioned above (Mixed).

**Supplementary Figure 2. Pipeline for the construction of different multiple sequence alignments used in this study.**

First, the full-length sequence is retrieved and used for MSA generation with MMseqs (see Methods). Then, delimited alignments are extracted by retrieving only columns corresponding to positions present in the 3D structure. For some predictions, peptide alignments are extended by up to 100 or 200 residue positions. Finally, three modes are used to combine evolutionary information of the receptor and peptide: a mixed alignment in which as many partner sequences as possible are paired while sequences with a single partner homolog present in a species are added as unpaired, a fully unpaired alignment in which no sequences are paired, and a mode with no evolutionary information for the peptide.

**Supplementary Figure 3. Prediction quality of individual test cases for different prediction modes.**

Illustrations of the DockQ score of the best predicted model (chosen as the best AF2 combined score) for each test case. Test cases are identified by their PDB code and sorted by the cumulative length of the input sequence (receptor+peptide). This length is indicated on top of each bar. PDB codes are color-coded according to the peptide bound conformation: helical (green), strand (orange), coil (purple) or a mixture (black). The dashed line indicates a DockQ score of 0.23, which is the threshold for an acceptable prediction as evaluated on protein-protein complexes (see Methods). Three prediction modes are used: **(a)** full-length partners, **(b)** delimited receptor and full-length ligand and **(c)** delimited receptor and ligand. All predictions in this figure are made with mixed alignments.

**Supplementary Figure 4. AlphaFold2-Multimer success rates on the benchmark dataset using ligands extended by fragments of size 100 and 200.**

Success rates as in main Figure 2, calculated for ligands extended by fragments of size 100 (for the two leftmost bars) and 200 (for the two rightmost bars) with either a mixed alignment or no peptide alignment.

**Supplementary Figure 5. Comparison of successful predictions in different prediction modes.**

UpSet diagram as in Figure 4a for different prediction modes: full-length with mixed alignment, delimited receptor with a ligand extended by a fragment of size 200 with mixed alignment, delimited receptor and peptide without MSA for the peptide, delimited receptor with a ligand extended by a fragment of size 100 with mixed alignment, delimited receptor and peptide with mixed alignment and delimited receptor and peptide with unpaired alignment.

**Supplementary Figure 6. Cross-partners AF2 predictions for complexes between receptors and cognate or non cognate peptides from the same cluster.**

Cross-partners analyses were performed between cases from the same groups of structures clustered according to the bound ligand conformations: **(a)** short helix, **(b)** medium helix, **(c)** long helix, **(d)** coil, **(e)** helix+strand, **(f)** single strand, **(g)** two strands. For each cross-partner configuration, three matrices are shown: DockQ scores, AF2 combined scores and percentage of overlap with correct interface residues. Matrix rows correspond to the same receptor (labeled as r.PDB) crossed with different peptides while columns report for every ligand (labeled as l.PDB) crossed with different receptors. Color code for DockQ score matrices is brown for Incorrect, orange for Acceptable, yellow for Medium and green for High; for AF2 confidence score matrices, the colors range from dark blue to light green for AF2 scores from 0 to 1; for the fraction of overlap with correct interface residues, the colors range from red to yellow for percentages between 0% to 100%. The cases represented as structural models in the panels (b), (c) and (d) of Figure 5 are labeled on the lines of the corresponding matrices with an asterisk and the corresponding panel letter.

**Supplementary Table 1**

### INDEX: Index of the case in benchmark dataset
### PDB: Reference of the experimental structure used for validation
### CHAIN: Label of the chain used in the reference structure
### REC/LIG: Status of the molecule either receptor or ligand. A case can have several receptor in case of homodimers or heterodimers
### UNIPROT: Reference sequence Uniprot index used to generate the multiple sequence alignments
### START: Index of the first residue modeled in the delimited conditions as defined in the SEQRES PDB parameter
### STOP: Index of the last residue modeled in the delimited conditions as defined in the SEQRES PDB parameter
### DELIM_LENG Length of the chain in the delimited condition
### FULL_LENGT Full length of the protein sequence as defined in Uniprot
### SS_TYPE: Secondary structures of ligand in the reference complex. Numbers in brackets report the length of each secondary struct. element

| #INDEX | PDB | CHAIN | REC/LIG | UNIPROT | START | STOP | DELIM_LENGT | FULL_LENGTH | SS_TYPE |
|---|---|---|---|---|---|---|---|---|---|
| 1 | 5NCL | A | receptor | P53894 | 251 | 756 | 506 | 756 | - |
| 1 | 5NCL | B | receptor | P43563 | 46 | 287 | 242 | 287 | - |
| 1 | 5NCL | D | ligand | P24276 | 205 | 214 | 10 | 1250 | coil |
| 2 | 5OJR | A | receptor | Q8DI95 | 39 | 347 | 309 | 347 | - |
| 2 | 5OJR | E | ligand | Q8DIV4 | 335 | 352 | 18 | 360 | strand (3) + strand (3) |
| 3 | 5V1U | A | receptor | D1CIZ5 | 1 | 88 | 88 | 88 | - |
| 3 | 5V1U | E | ligand | D1CIY7 | 1 | 20 | 20 | 65 | strand (2) + strand (6) |
| 4 | 6A30 | A | receptor | Q62768 | 944 | 1523 | 580 | 1735 | - |
| 4 | 6A30 | P | ligand | P63045 | 87 | 92 | 6 | 116 | coil |
| 5 | 6DO3 | A | receptor | Q9Y2U9 | 1 | 362 | 362 | 406 | - |
| 5 | 6DO3 | C | ligand | Q9Y6D0 | 85 | 91 | 7 | 93 | helical (5) |
| 6 | 6G04 | A | receptor | Q06672 | 1 | 152 | 152 | 205 | - |
| 6 | 6G04 | B | ligand | P39938 | 100 | 119 | 20 | 119 | helix (6) + coil |
| 7 | 6ICV | A | receptor | Q86TU7 | 1 | 503 | 503 | 594 | - |
| 7 | 6ICV | C | ligand | P60709 | 66 | 88 | 23 | 375 | coil |
| 8 | 6IDX | A | receptor | Q96JJ3 | 5 | 515 | 511 | 720 | - |
| 8 | 6IDX | C | ligand | Q3UHD1 | 1471 | 1495 | 25 | 1582 | helix (16) |
| 9 | 6IXQ | A | receptor | P19524 | 1152 | 1574 | 423 | 1574 | - |
| 9 | 6IXQ | B | ligand | P32364 | 615 | 650 | 36 | 656 | strand (2) |
| 10 | 6J0W | A | receptor | P38850 | 1 | 513 | 513 | 1070 | - |
| 10 | 6J0W | C | ligand | P40026 | 13 | 41 | 29 | 464 | strand (3) |
| 11 | 6J0X | A | receptor | P38850 | 1 | 513 | 513 | 1070 | - |
| 11 | 6J0X | E | ligand | Q06164 | 22 | 37 | 16 | 1454 | strand (3) + helix (5) |
| 12 | 6JLH | A | receptor | Q5XJX1 | 7 | 267 | 261 | 272 | - |
| 12 | 6JLH | B | ligand | P60880 | 154 | 170 | 17 | 206 | helix (16) |
| 13 | 6JWJ | A | receptor | P33755 | 113 | 580 | 468 | 580 | - |
| 13 | 6JWJ | C | ligand | P53044 | 288 | 305 | 18 | 361 | strand (3) + coil |
| 14 | 6KPB | C | receptor | Q9LPR8 | 1 | 482 | 482 | 482 | - |
| 14 | 6KPB | B | ligand | Q700D2 | 367 | 383 | 17 | 503 | helix (13) |
| 15 | 6L0V | A | receptor | F4K0X5 | 1006 | 1066 | 61 | 1075 | - |
| 15 | 6L0V | B | ligand | Q5XVG3 | 274 | 287 | 14 | 287 | strand (3) + strand (5) |
| 16 | 6LPH | A | receptor | Q9VG38 | 1 | 258 | 258 | 468 | - |
| 16 | 6LPH | B | ligand | P23647 | 363 | 387 | 25 | 805 | strand (5) + helix (5) |
| 17 | 6OCG | A | receptor | Q7L8A9 | 59 | 305 | 247 | 365 | - |
| 17 | 6OCG | B | ligand | Q8N300 | 26 | 51 | 26 | 66 | helix (25) |
| 18 | 6PSD | G | receptor | Q9BSW2 | 47 | 121 | 75 | 731 | - |
| 18 | 6PSD | H | ligand | B3KM42 | 287 | 311 | 25 | 377 | helix (10) |
| 19 | 6PSE | A | receptor | Q8TD16 | 1 | 98 | 98 | 824 | - |
| 19 | 6PSE | B | receptor | Q8TD16 | 1 | 98 | 98 | 824 | - |
| 19 | 6PSE | C | ligand | Q9Y6G9 | 433 | 458 | 26 | 523 | helix (10) |
| 20 | 6RKO | A | receptor | P0ABJ9 | 1 | 522 | 522 | 522 | - |
| 20 | 6RKO | H | ligand | A5A618 | 1 | 29 | 29 | 29 | helix (25) |
| 21 | 6RKO | A | receptor | P0ABJ9 | 1 | 522 | 522 | 522 | - |
| 21 | 6RKO | X | ligand | P56100 | 1 | 37 | 37 | 37 | helix (24) |
| 22 | 6SAT | A | receptor | Q8NNN6 | 64 | 152 | 89 | 152 | - |
| 22 | 6SAT | B | receptor | Q8NNN6 | 64 | 152 | 89 | 152 | - |
| 22 | 6SAT | P | ligand | P94337 | 433 | 442 | 10 | 442 | helix (5) |
| 23 | 6TWN | B | receptor | P26039 | 1359 | 1659 | 301 | 2541 | - |
| 23 | 6TWN | C | ligand | P06493 | 207 | 223 | 17 | 297 | helix (14) |
| 24 | 6XFK | A | receptor | P0CL43 | 84 | 147 | 64 | 147 | - |
| 24 | 6XFK | B | ligand | P35672 | 543 | 558 | 16 | 562 | helix (10) |
| 25 | 6YN0 | A | receptor | P02919 | 58 | 804 | 747 | 844 | - |
| 25 | 6YN0 | B | ligand | P29131 | 75 | 93 | 19 | 319 | helix (12) |
| 26 | 7B1J | A | receptor | Q9Y6D9 | 597 | 718 | 122 | 718 | - |
| 26 | 7B1J | B | receptor | Q9Y6D9 | 597 | 718 | 122 | 718 | - |
| 26 | 7B1J | C | ligand | O43683 | 455 | 479 | 25 | 1085 | helix (15) |
| 27 | 7CFC | A | receptor | A1ZAC4 | 272 | 512 | 241 | 746 | - |
| 27 | 7CFC | F | ligand | Q7PLK0 | 63 | 78 | 16 | 867 | helix (9) |
| 28 | 7CZM | A | receptor | Q8TDY2 | 1490 | 1594 | 105 | 1594 | - |
| 28 | 7CZM | C | ligand | Q96CV9 | 173 | 185 | 13 | 577 | strand (3) |
| 29 | 7F2D | A | receptor | P93026 | 20 | 182 | 163 | 623 | - |
| 29 | 7F2D | B | ligand | P15455 | 468 | 472 | 5 | 472 | strand (2) |
| 30 | 7MKK | B | receptor | Q9W3W6 | 14 | 90 | 77 | 3313 | - |
| 30 | 7MKK | C | ligand | Q9W2H9 | 83 | 109 | 27 | 541 | helix (18) |

| | | | | | | | | | |
|---|---|---|---|---|---|---|---|---|---|
| 31 | 7MU2 | A | receptor | Q9Y4P8 | 11 | 363 | 353 | 454 | - |
| 31 | 7MU2 | B | ligand | E7EVC7 | 207 | 230 | 24 | 624 | helix (19) |
| 32 | 7NW1 | AAA | receptor | Q9Y3C8 | 1 | 167 | 167 | 167 | - |
| 32 | 7NW1 | FFF | ligand | Q9GZZ9 | 389 | 404 | 16 | 404 | helix (11) |
| 33 | 7QDW | A | receptor | Q8I2Y4 | 265 | 332 | 68 | 332 | - |
| 33 | 7QDW | B | ligand | Q8IK99 | 817 | 841 | 25 | 852 | helix (18) |
| 34 | 7RXQ | A | receptor | Q9BR39 | 1 | 437 | 437 | 696 | - |
| 34 | 7RXQ | B | ligand | P07293 | 1594 | 1609 | 16 | 1873 | coil |
| 35 | 7SID | A | receptor | Q13315 | 1 | 3056 | 3056 | 3056 | - |
| 35 | 7SID | B | ligand | O60934 | 727 | 754 | 28 | 754 | coil |
| 36 | 6J08 | A | receptor | Q3KP22 | 2 | 109 | 108 | 176 | - |
| 36 | 6J08 | D | ligand | Q8NHR7 | 174 | 209 | 36 | 220 | strand (2) + strand (3) + strand (5) |
| 37 | 6ZW0 | A | receptor | Q57968 | 2 | 293 | 292 | 293 | - |
| 37 | 6ZW0 | C | ligand | Q58261 | 140 | 169 | 30 | 241 | helix (8) + strand (3) + strand (4) |
| 38 | 7N40 | A | receptor | Q09028 | 1 | 425 | 425 | 425 | - |
| 38 | 7N40 | B | receptor | Q5TKA1 | 98 | 274 | 177 | 542 | - |
| 38 | 7N40 | C | ligand | Q96GY3 | 95 | 126 | 32 | 246 | strand (4) + strand (4) + helix (3) + helix (11) |
| 39 | 7O6N | A | receptor | Q20057 | 1 | 99 | 99 | 113 | - |
| 39 | 7O6N | D | ligand | O76616 | 179 | 193 | 15 | 307 | helix (14) |
| 40 | 5OW5 | A | receptor | Q8BG40 | 481 | 658 | 178 | 658 | - |
| 40 | 5OW5 | B | receptor | E9PZI6 | 3 | 80 | 78 | 493 | - |
| 40 | 5OW5 | E | ligand | Q80VC9 | 461 | 470 | 10 | 1252 | helix (9) |
| 41 | 6JMT | A | receptor | Q80XR8 | 1 | 360 | 360 | 679 | - |
| 41 | 6JMT | L | ligand | Q9ES28 | 685 | 705 | 21 | 862 | helix (14) |
| 42 | 6GP7 | B | receptor | P0CI74 | 4 | 64 | 61 | 98 | - |
| 42 | 6GP7 | C | receptor | P0CI74 | 4 | 64 | 61 | 98 | - |
| 42 | 6GP7 | D | ligand | A0A5D4ND23 | 1 | 17 | 17 | 908 | helix (9) |

**Supplementary Table 2**

**# Interfaces involving the folding of a short-length helix ligand (5-6 residues)**

| index | pdb |
|---|---|
| 5 | 6DO3 |
| 6 | 6G04 |

**# Interfaces involving the folding of a medium-length helix ligand (9-11 residues)**

| | |
|---|---|
| 18 | 6PSD |
| 24 | 6XFK |
| 27 | 7CFC |
| 32 | 7NW1 |

**# Interfaces involving the folding of a long-length helix ligand (12-16 residues)**

| | |
|---|---|
| 8 | 6IDX |
| 14 | 6KPB |
| 23 | 6TWN |
| 25 | 6YN0 |

**# Interfaces involving the folding of a coil ligand**

| | |
|---|---|
| 4 | 6A30 |
| 7 | 6ICV |
| 34 | 7RXQ |

**# Interfaces involving the folding of a helix + strand ligand**

| | |
|---|---|
| 11 | 6J0X |
| 16 | 6LPH |

**# Interfaces involving the folding of a single strand ligand (2-3 residues)**

| | |
|---|---|
| 9 | 6IXQ |
| 10 | 6J0W |
| 13 | 6JWJ |
| 28 | 7CZM |
| 29 | 7F2D |

**# Interfaces involving the folding of a two-stranded ligand**

| | |
|---|---|
| 2 | 5OJR |
| 3 | 5V1U |
| 15 | 6L0V |

**Supp. Figure 1**

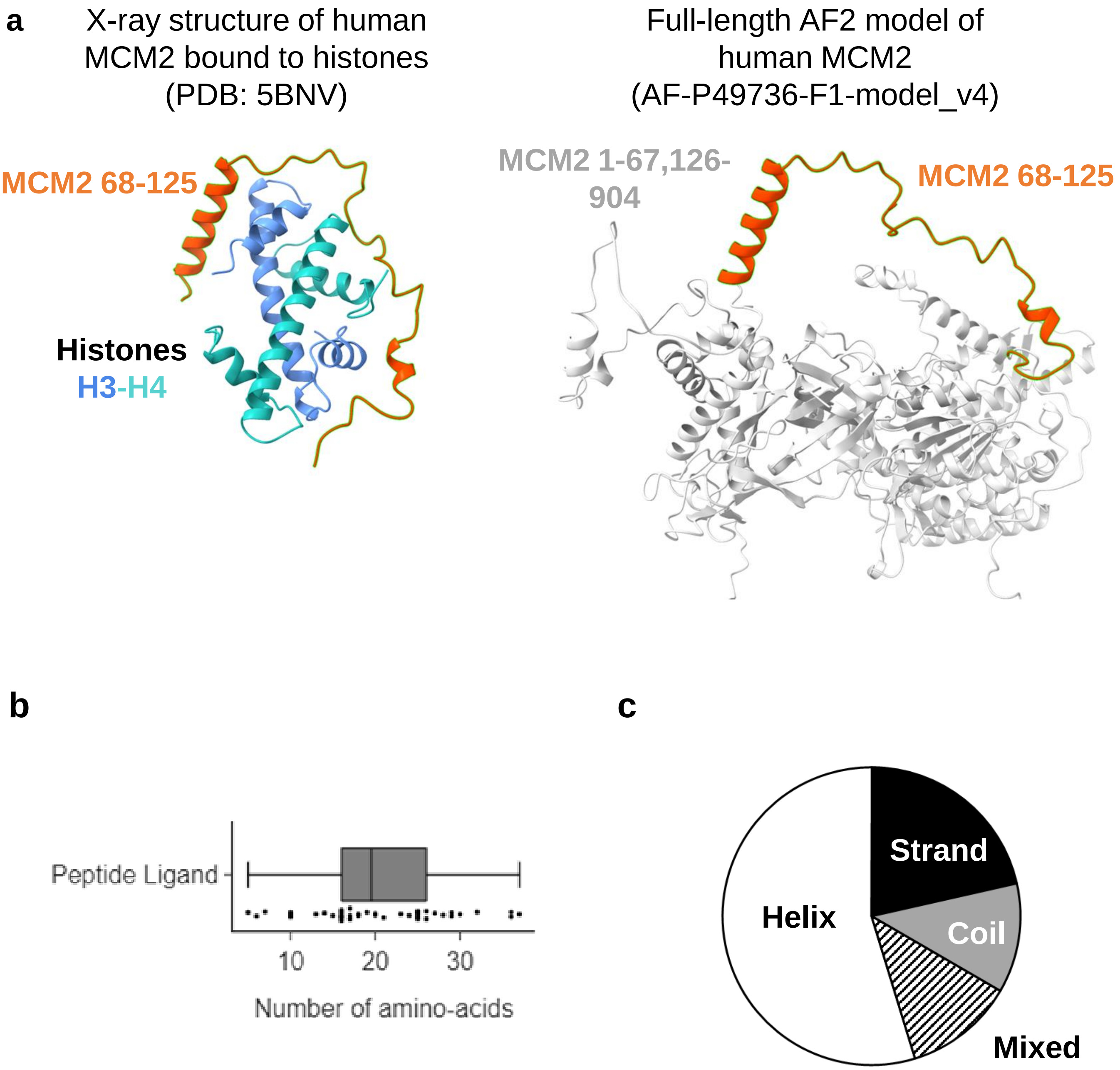

**Supp. Figure 2**

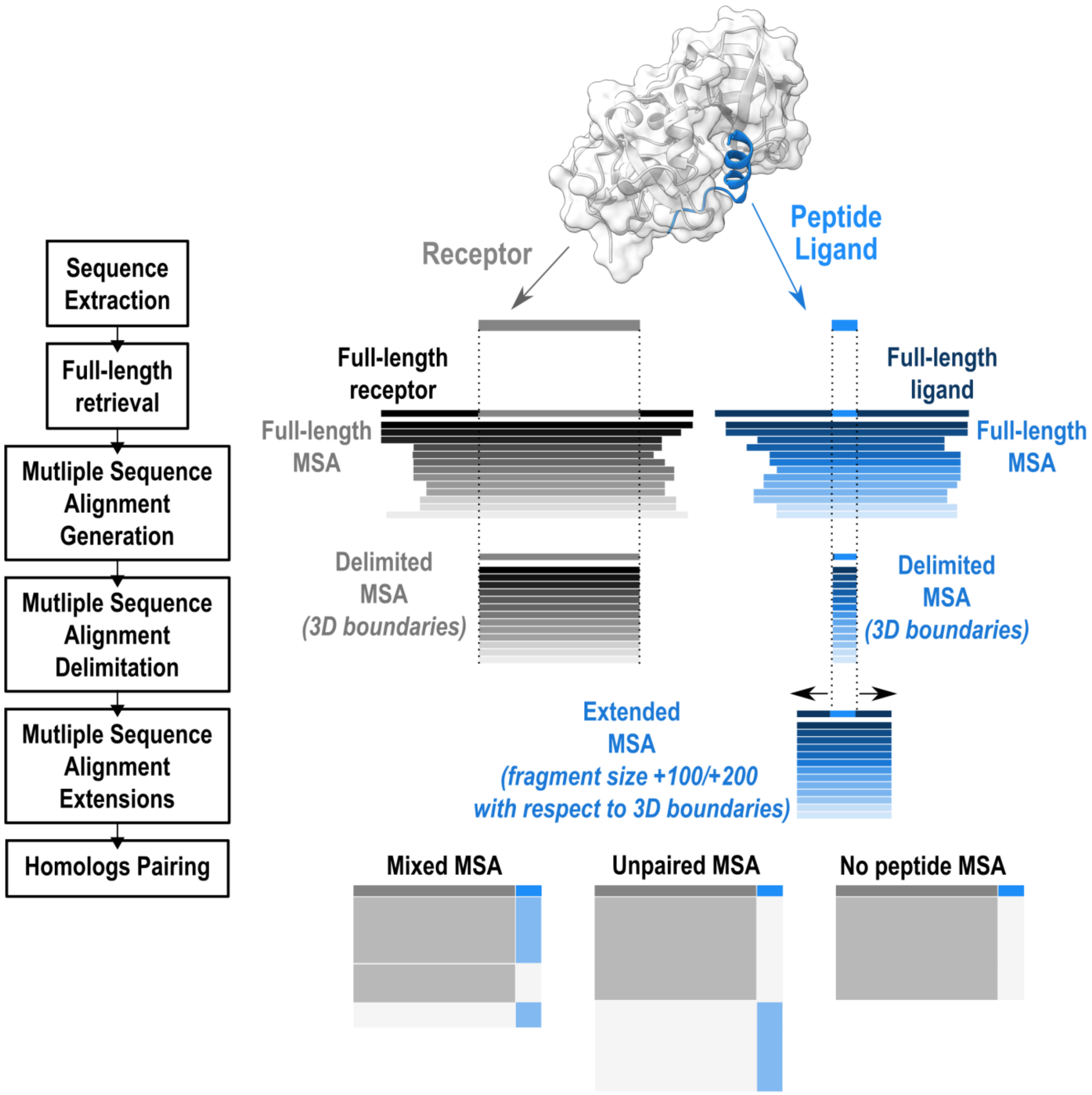

#### Supp. Figure 3

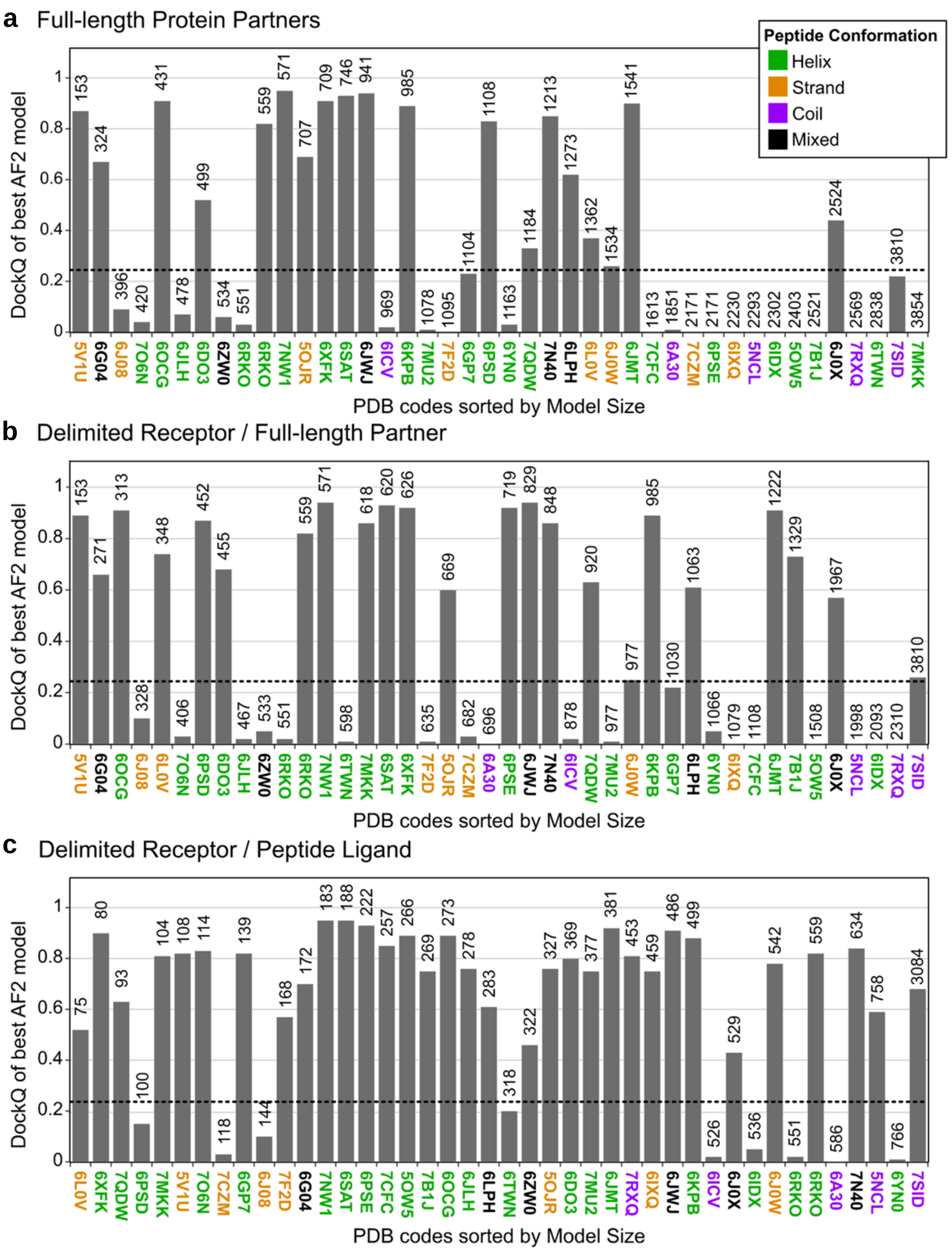

**Supp. Figure 4**

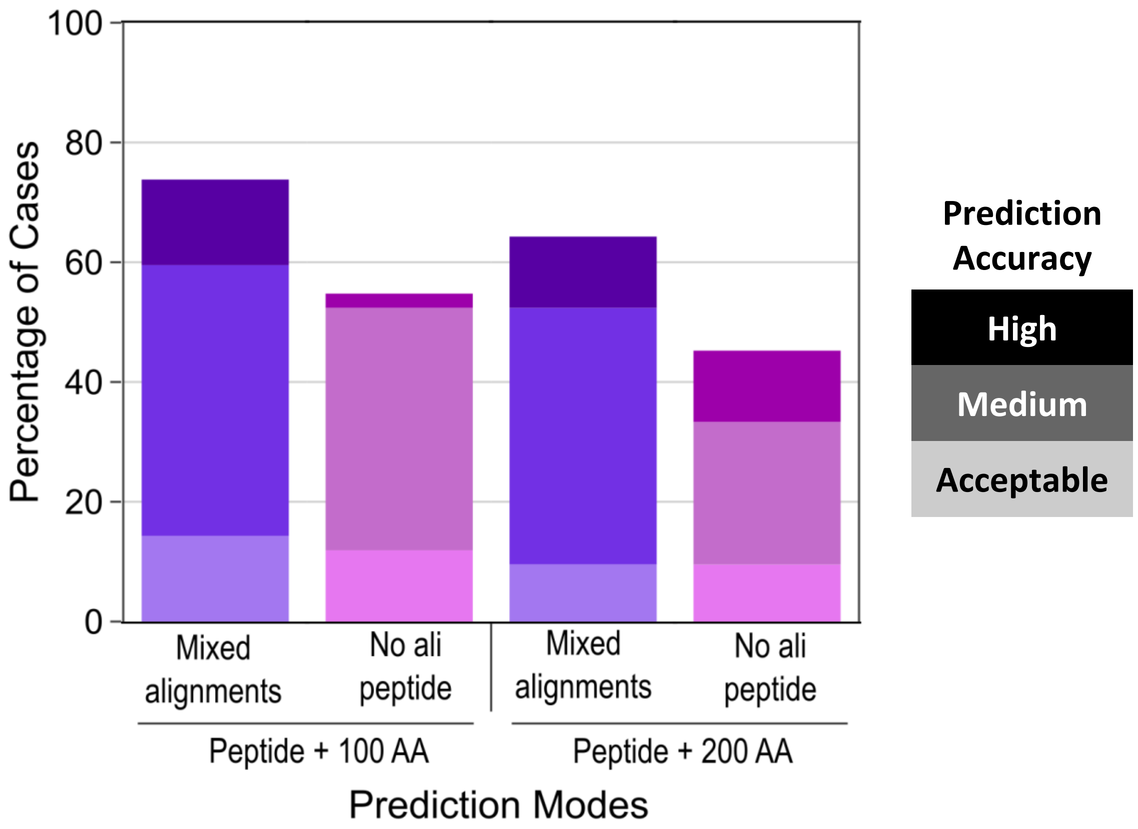

#### Supp. Figure 5

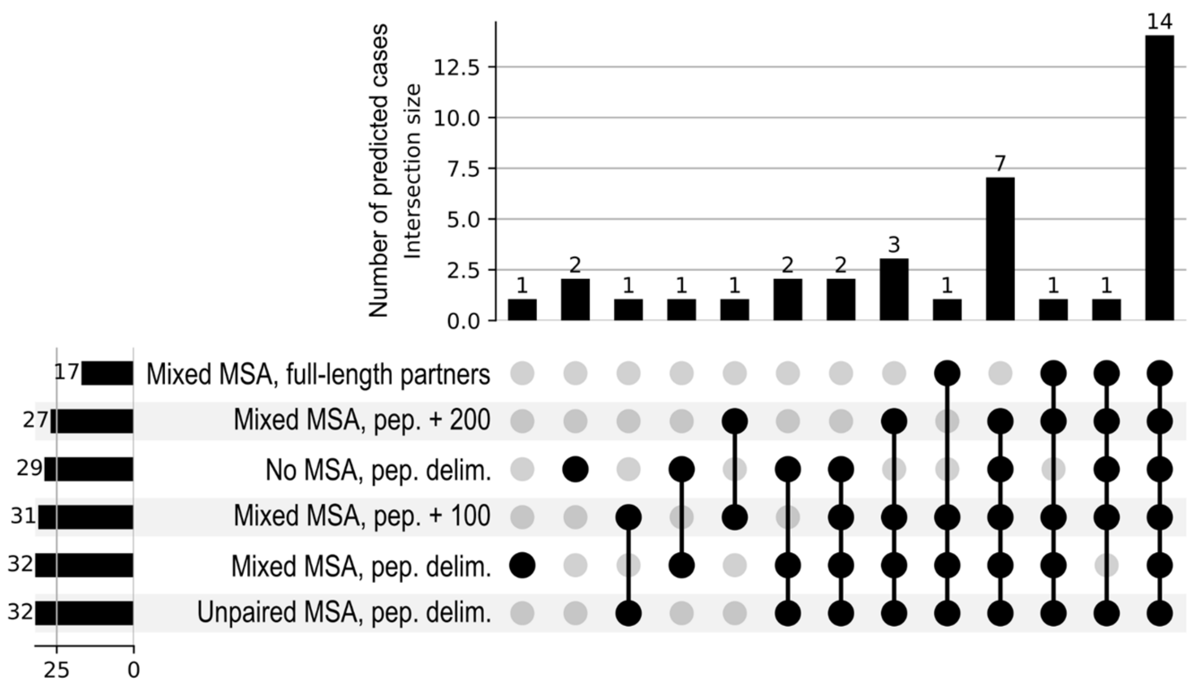

**Supp. Figure 6**

*a. Interfaces involving the folding of a short-length helix ligand (5-6 res.)*

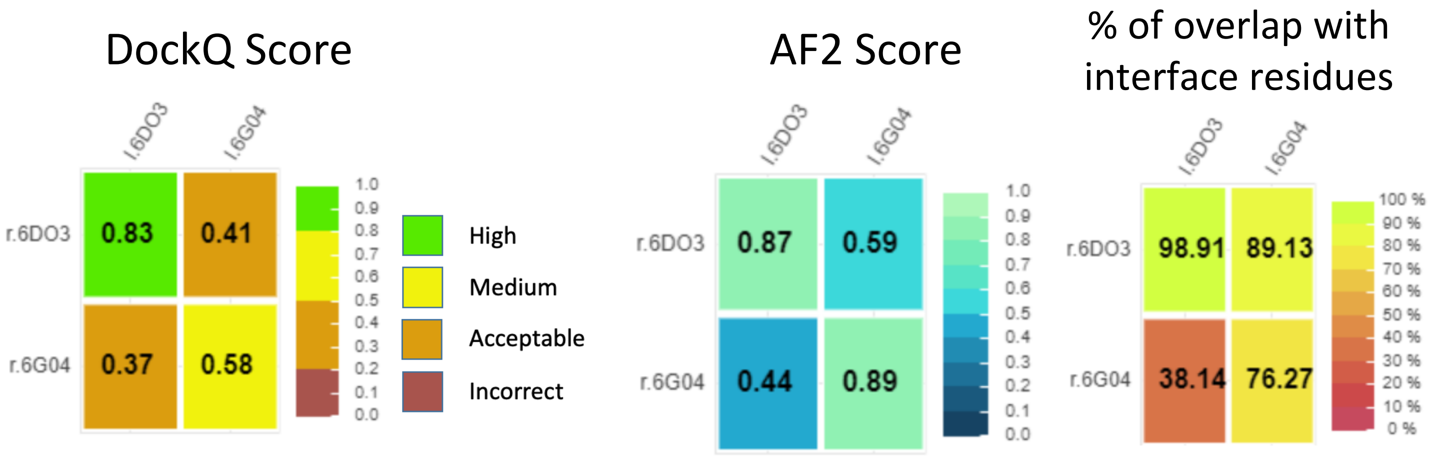


*b. Interfaces involving the folding of a medium-length helix ligand (9-11 res.)*

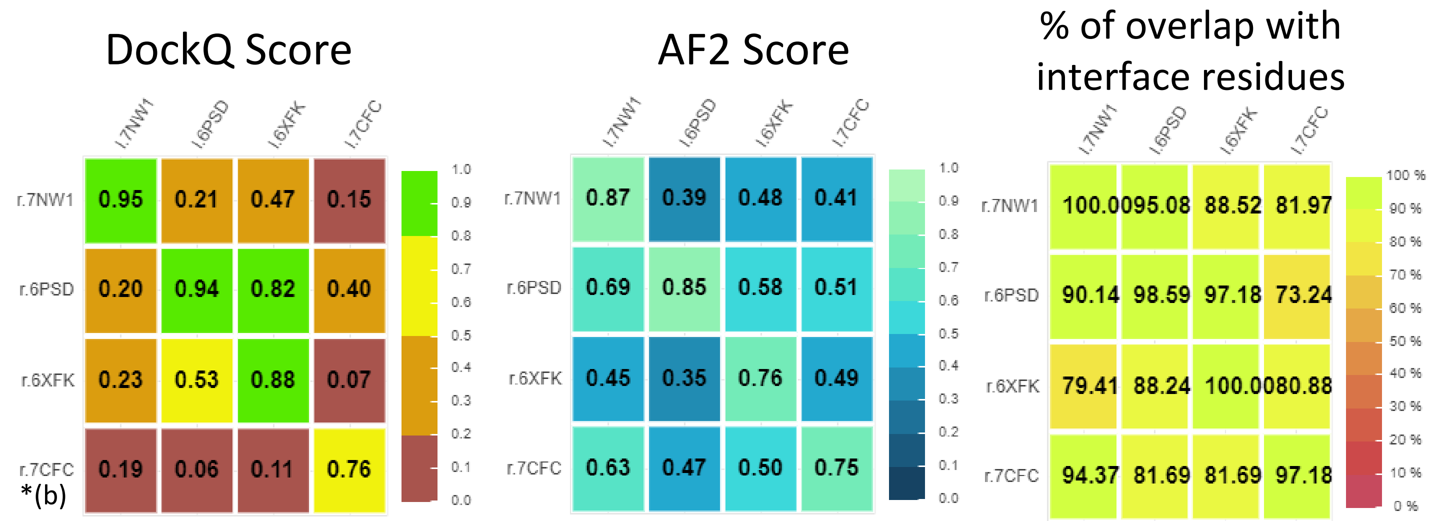


*c. Interfaces involving the folding of a long-length helix ligand (12-16 res.)*

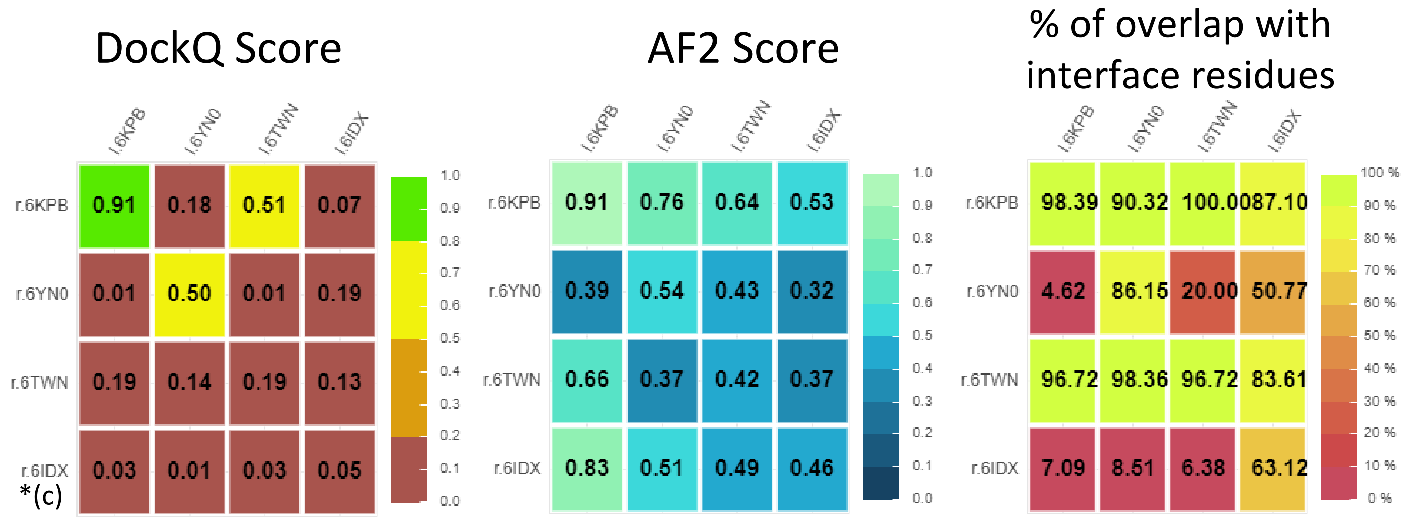

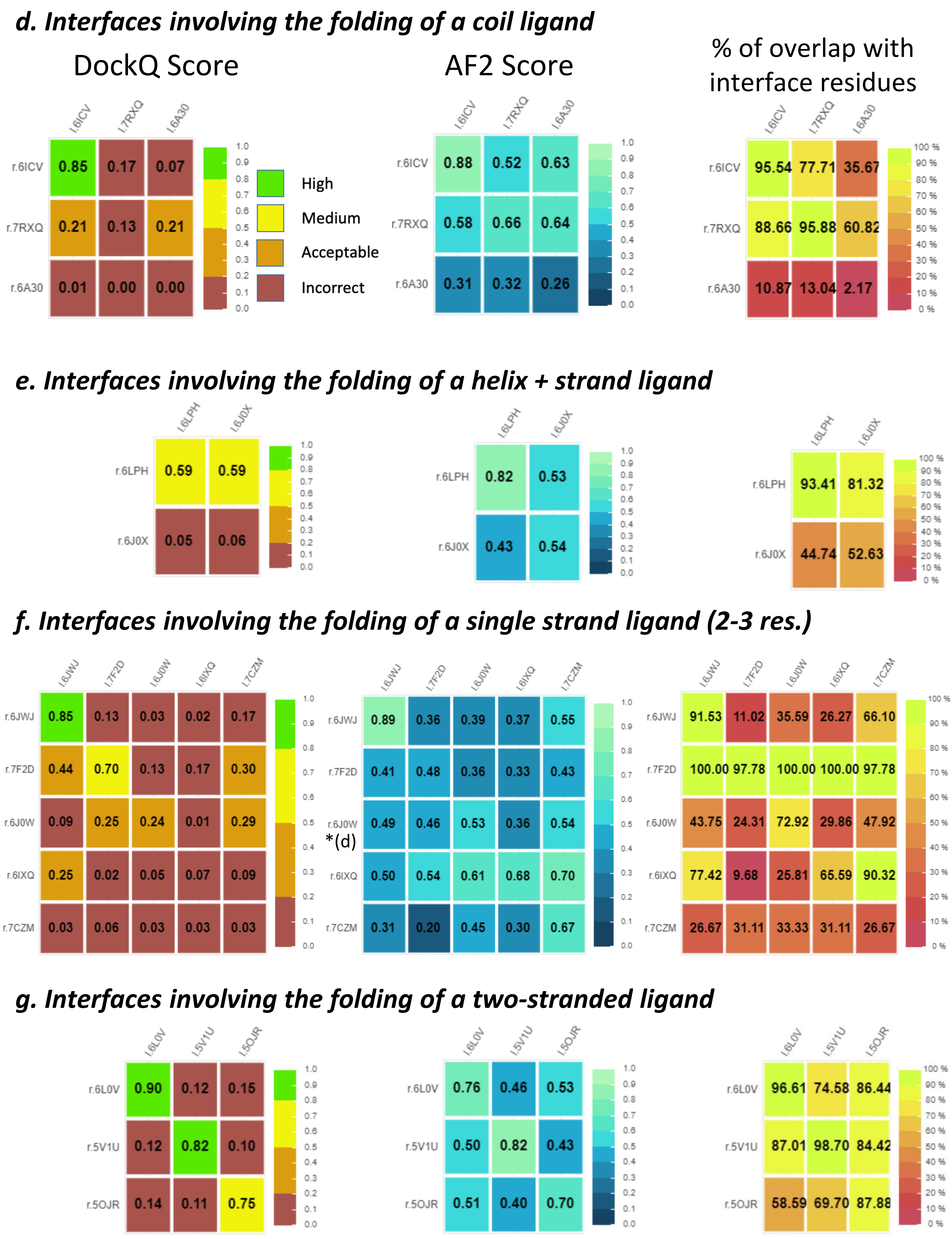
d. Interfaces involving the folding of a coil ligand
DockQ Score
AF2 Score
% of overlap with interface residues
l.6ICV l.7RXQ l.6A30
r.6ICV 0.85 0.17 0.07
r.7RXQ 0.21 0.13 0.21
r.6A30 0.01 0.00 0.00
High
Medium
Acceptable
Incorrect
r.6ICV 0.88 0.52 0.63
r.7RXQ 0.58 0.66 0.64
r.6A30 0.31 0.32 0.26
r.6ICV 95.54 77.71 35.67
r.7RXQ 88.66 95.88 60.82
r.6A30 10.87 13.04 2.17
e. Interfaces involving the folding of a helix + strand ligand
l.6LPH l.6J0X
r.6LPH 0.59 0.59
r.6J0X 0.05 0.06
r.6LPH 0.82 0.53
r.6J0X 0.43 0.54
r.6LPH 93.41 81.32
r.6J0X 44.74 52.63
f. Interfaces involving the folding of a single strand ligand (2-3 res.)
l.6JWJ l.7F2D l.6J0W l.6IXQ l.7CZM
r.6JWJ 0.85 0.13 0.03 0.02 0.17
r.7F2D 0.44 0.70 0.13 0.17 0.30
r.6J0W 0.09 0.25 0.24 0.01 0.29
r.6IXQ 0.25 0.02 0.05 0.07 0.09
r.7CZM 0.03 0.06 0.03 0.03 0.03
r.6JWJ 0.89 0.36 0.39 0.37 0.55
r.7F2D 0.41 0.48 0.36 0.33 0.43
r.6J0W 0.49 0.46 0.53 0.36 0.54
*(d)
r.6IXQ 0.50 0.54 0.61 0.68 0.70
r.7CZM 0.31 0.20 0.45 0.30 0.67
r.6JWJ 91.53 11.02 35.59 26.27 66.10
r.7F2D 100.00 97.78 100.00 100.00 97.78
r.6J0W 43.75 24.31 72.92 29.86 47.92
r.6IXQ 77.42 9.68 25.81 65.59 90.32
r.7CZM 26.67 31.11 33.33 31.11 26.67
g. Interfaces involving the folding of a two-stranded ligand
l.6L0V l.5V1U l.5OJR
r.6L0V 0.90 0.12 0.15
r.5V1U 0.12 0.82 0.10
r.5OJR 0.14 0.11 0.75
r.6L0V 0.76 0.46 0.53
r.5V1U 0.50 0.82 0.43
r.5OJR 0.51 0.40 0.70
r.6L0V 96.61 74.58 86.44
r.5V1U 87.01 98.70 84.42
r.5OJR 58.59 69.70 87.88
